# Rewiring of Integrin Signaling and Cell-cycle Deregulation Drive SMARCB1-Deficient Epithelioid Sarcoma

**DOI:** 10.64898/2026.09.01.748640

**Authors:** Ryo Miyamoto, Jiyeon Park, Gabrielle H Dwosh, Valentina Zuco, Sandro Pasquali, Brendan C Dickson, David G Kirsch

**Author notes:** These authors contributed equally (Ryo Miyamoto, Jiyeon Park).

## Abstract

Epithelioid sarcoma (EPS) is an aggressive soft-tissue sarcoma characterized by loss of the chromatin-remodeling subunit SMARCB1. The oncogenic programs driving EPS remain poorly understood. Through CRISPR loss-of-function screens, we identified conserved dependencies on integrin signaling components and cyclin-dependent kinases (CDKs). Genetic disruption of integrin subunit alpha V (ITGAV)-centered signaling impaired epithelioid cluster formation and reduced MYC expression. SMARCB1 re-expression phenocopied these effects and revealed that SMARCB1 loss selectively represses context-dependent integrin subunits while preserving an ITGAV-centered pro-survival axis, associated with altered BAF complex occupancy. Analysis of EPS cell lines and primary tumors revealed frequent genetic or epigenetic inactivation of CDKN2A/p16, indicating that loss of cell-cycle control is a key cooperating event in EPS development and providing a mechanistic rationale for targeting CDK4/6. Together, these findings establish integrin-driven oncogenic signaling coupled with disruption of cell-cycle control as a central oncogenic program in EPS and identify actionable therapeutic vulnerabilities.

## Introduction

Epithelioid sarcoma (EPS) is a soft-tissue sarcoma that primarily affects the distal extremities of adolescents and young adults^1,2^. EPS was first described by Enzinger in 1970 as a distinct sarcoma with epithelial morphology despite its presumed mesenchymal origin^3^. EPS accounts for approximately 1% of all sarcomas and has a poor prognosis, with a 5-year survival rate of approximately 50% due to frequent local recurrence and distal metastasis^1,2^.

EPS is immunohistochemically characterized by loss of SMARCB1, a core subunit of the BAF (SWI/SNF) chromatin remodeling complex. The BAF complex consists of 10–15 subunits, with multiple distinct assemblies characterized to date, and regulates gene expression through ATP-dependent nucleosome remodeling and dynamic control of chromatin accessibility^4,5^. Genes encoding BAF subunits are mutated in nearly 25% of cancers, underscoring the tumor-suppressive role of this complex. Despite genomic complexity, protein loss of SMARCB1 is observed in approximately 90% of EPS cases^2,6^. Loss of SMARCA4, another BAF component, accounts for a portion of the remaining cases, indicating the central role of BAF complex dysfunction in the pathogenesis of EPS^7,8^. However, the oncogenic programs resulting from SMARCB1 loss remain poorly understood. The rarity and biological heterogeneity of EPS have hindered systematic functional studies. Such studies could define key oncogenic programs resulting from BAF dysfunction and reveal exploitable therapeutic vulnerabilities in EPS.

SMARCB1 deficiency is also a hallmark of rhabdoid tumors (RTs), which are aggressive cancers affecting infants and children^9^. RTs exhibit an exceptionally low mutational burden, with biallelic SMARCB1 inactivation serving as the principal oncogenic driver. This genetic simplicity has made RTs the dominant experimental model to study the consequences of SMARCB1 inactivation, often by restoring the expression of SMARCB1. In RTs, loss of SMARCB1 impairs BAF complex integrity and occupancy at enhancers^10,11^. This results in widespread epigenetic reprogramming associated with functional antagonism between the BAF complex and the polycomb-repressive complex 2 (PRC2), leading to loss of active enhancer marks such as H3K27ac, accumulation of the repressive mark H3K27me3, and ultimately transcriptional silencing of BAF target genes. A key downstream consequence is the repression of cyclin-dependent kinase (CDK) inhibitors, including CDKN2A (p16^INK4A^) and CDKN1A (p21^CIP1^), leading to increased CDK activity and accelerated cell cycle progression^12,13^.

Insights from RTs have informed therapeutic strategies for EPS, most notably tazemetostat, a selective EZH2 inhibitor that received accelerated approval for patients with advanced EPS^14,15^. However, clinical responses to EZH2 inhibition have been limited, with an objective response rate of approximately 15%. EPS differs from RT in anatomical distribution, histological heterogeneity, age of onset, mutational burden, and potentially distinct cells of origin. These differences raise questions about the generalizability of RT-derived therapeutic paradigms to EPS. More recently, secondary hematologic malignancies were reported in a clinical trial of tazemetostat in patients with follicular lymphoma, leading to voluntary withdrawal from all indications, including EPS.^16^ These clinical challenges highlight the unmet medical need and emphasize the importance of EPS-specific functional studies.

To systematically define cancer dependencies in EPS, we performed CRISPR loss-of-function screens focused on druggable genes across a panel of human EPS cell lines. These screens revealed a lack of dependency on EZH2, whereas EPS cell lines exhibited consistent dependencies on integrin signaling components and CDKs. Here, we characterize mechanisms underlying these dependencies and their contribution to the malignant phenotype of EPS.

## Results

### Druggable CRISPR screens reveal dependencies and actionable drug targets in EPS

To characterize EPS dependencies, we first analyzed DepMap CRISPR screening data, which includes two SMARCB1-deficient EPS cell lines, CCLFPEDS0008T (CCLFP8) and VAESBJ (VAES). Genes with dependency scores at least 0.2 below the pan-cancer average were extracted to define shared and cell line-specific dependencies and associated pathway enrichment (**Fig. 1A**). Shared dependencies included the BAF complex subunit (SMARCD1), integrin signaling genes (ITGAV, RAC1, CDC42, ELMO2, PTPN11, GRB2), and endocytosis mediators (STAMBP, CHMP4B). Additional BAF components, including SMARCA4 and SMARCC2, were identified as cell line-specific dependencies, suggesting synthetic lethality associated with BAF complex perturbation. The integrin-associated adaptor proteins CRK and CRKL were selectively essential in VAES and CCLFP8, respectively, further supporting a pathway-level dependency on integrin signaling.

**Fig. 1.**
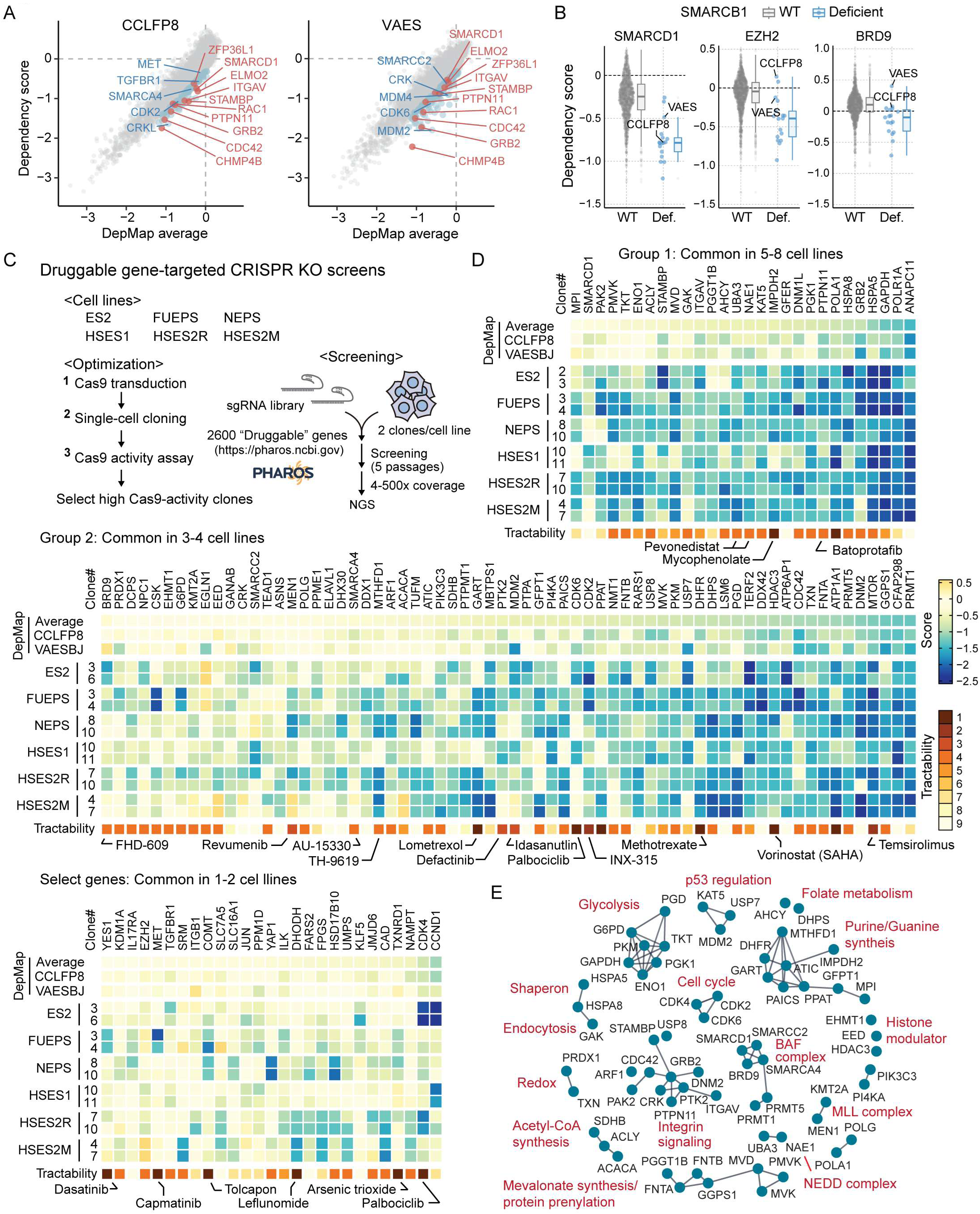
| Druggable gene-focused CRISPR screens identify core EPS vulnerabilities (A) DepMap dependency scores for two EPS cell lines (CCLFP8 and VAES) across 17,917 genes plotted against pan- cancer average scores (from 1,178 cell lines). Dependency genes specific to each EPS cell line or shared by both cell lines are indicated in blue and red, respectively. (B) Two-class comparison of dependency scores between SMARCB1 wild-type (WT; n = 1,161) and SMARCB1-deficient cell lines (n = 17) for SMARCD1, BRD9 and EZH2. (C) Schematic of the druggable gene-focused CRISPR screen. Druggable genes were selected from the Pharos database. (D) Heatmaps of CRISPR-screen dropout scores across six EPS cell lines, each analyzed using two independent single- cell clones. The DepMap pan-cancer average and DepMap scores for CCLFP8 and VAES are displayed alongside for comparison. Screen hits were identified in each EPS cell line and classified by hit frequency across eight lines, including CCLFP8 and VAES: Group 1 (5–8 cell lines), Group 2 (3–4), or Group 3 (1–2 cell lines). For visualization, Group 3 genes were further filtered based on druggability or reported relevance to cancer biology. (E) Network plot of pathway enrichment among screen hits. Group 1 and 2 genes were analyzed for physical and/or functional interactions using STRING. CDK4, a Group 3 gene, was also included because of its mutually exclusive dependency pattern between CDK6.

We next extended our analysis to SMARCB1-deficient cancer cell lines in DepMap, including EPS, RT, and atypical teratoid rhabdoid tumor (ATRT) models. SMARCD1 dependency was shared across SMARCB1- deficient lines (**Fig. 1B**). In contrast, BRD9 and EZH2, previously proposed synthetic-lethal targets in BAF-perturbed cancers, were dispensable in EPS cell lines, reinforcing biological differences between EPS and RT.

To systematically map EPS fitness genes beyond the two EPS cell lines included in DepMap (CCLFP8 and VAES), we performed a focused CRISPR knockout (KO) screen in six additional human EPS cell lines (ES2, FUEPS, HSES1, HSES2R, HSES2M, and NEPS) (**Supplementary Fig. 1A, Supplementary Table 1**). The sgRNA library targeted 2,600 druggable genes and included BAF components and related genes (SMARCA1 and SMARCA5 from the ISWI complex) regardless of druggability. For each cell line, we stably expressed Cas9, established single-cell-derived clones, and selected two independent clones with high Cas9 activity per cell line to maximize editing efficacy while minimizing the chance of capturing clone-specific phenotypes (**Fig. 1C, Supplementary Fig. 1B**). Correlation between replicate scores, together with the expected enrichment of canonical tumor suppressors (TP53, CDKN1A) and depletion of core essential genes (PLK1, RPA1), indicated robust screen performance (**Supplementary Fig. 1C, Supplementary Data 1**).

To identify EPS dependencies, we normalized depletion scores to the DepMap scale and stratified genes by hit frequency across eight EPS cell lines, including six screened and two cell lines from DepMap (Group 1, 5–8 lines; Group 2, 3–4; Group 3, 1–2). Gene tractability scores were incorporated for target prioritization^17^.

Consistent with the prior analysis of CCLFP8 and VAES, Group 1 included STAMBP, ITGAV and PTPN11. Metabolic enzymes such as PMVK, MVD, TKT and ENO1 also fell into the same group (**Fig. 1D**). Group 2 highlighted intrinsic heterogeneity among EPS models: three lines were dependent on BRD9, whereas five were resistant. Other BAF components, including ARID2, SMARCC2, and SMARCA4, showed similar cell line-specific dependency patterns (**Fig. 1D**, **Supplementary Fig. 1D**). CDK4 and CDK6 showed mutually exclusive dependencies across EPS lines, indicating a conserved requirement for CDK-driven cell-cycle progression. By contrast, EZH2 KO had minimal effects, and receptor tyrosine kinases, which are within a highly tractable gene category, were broadly nonessential (**Supplementary Fig. 1D**).

Protein-protein interaction network analysis of screen hits showed enrichment of integrin signaling, BAF complex components, and CDKs (**Fig. 1E**). Glycolysis, nucleotide biosynthesis, mevalonate/prenylation metabolism were also overrepresented, suggesting metabolic rewiring in EPS.

### ITGAV-centered integrin signaling is associated with the epithelial-like phenotype of EPS

The enrichment of integrin signaling genes, together with the central role of integrins in epithelial-mesenchymal plasticity, prompted us to examine their functional contribution to EPS fitness. Pan-cancer DepMap analysis showed correlated dependencies between ITGAV and downstream mediators of integrin signaling, including PTK2 (FAK), ELMO2 and RAC1: this pattern was preserved in EPS cell lines (**Fig. 2A, B**). CRISPR knockout in an extended panel of EPS lines confirmed that ITGAV is required for EPS cell adhesion and proliferation (**Fig. 2C, D**, **Supplementary Fig. 2A–D**). Other integrin components also showed strong fitness dependencies, with ITGB1, CRK and CRKL representing cell line-specific core essentials, suggestive of divergent downstream wiring across EPS models. Notably, growth inhibition was accompanied by loss of clustered morphology characteristic of EPS and occasional emergence of elongated, process-bearing cells with reduced cell-cell adhesion (**Fig. 2D**, **Supplementary Fig. 2A–D**).

**Fig. 2.**
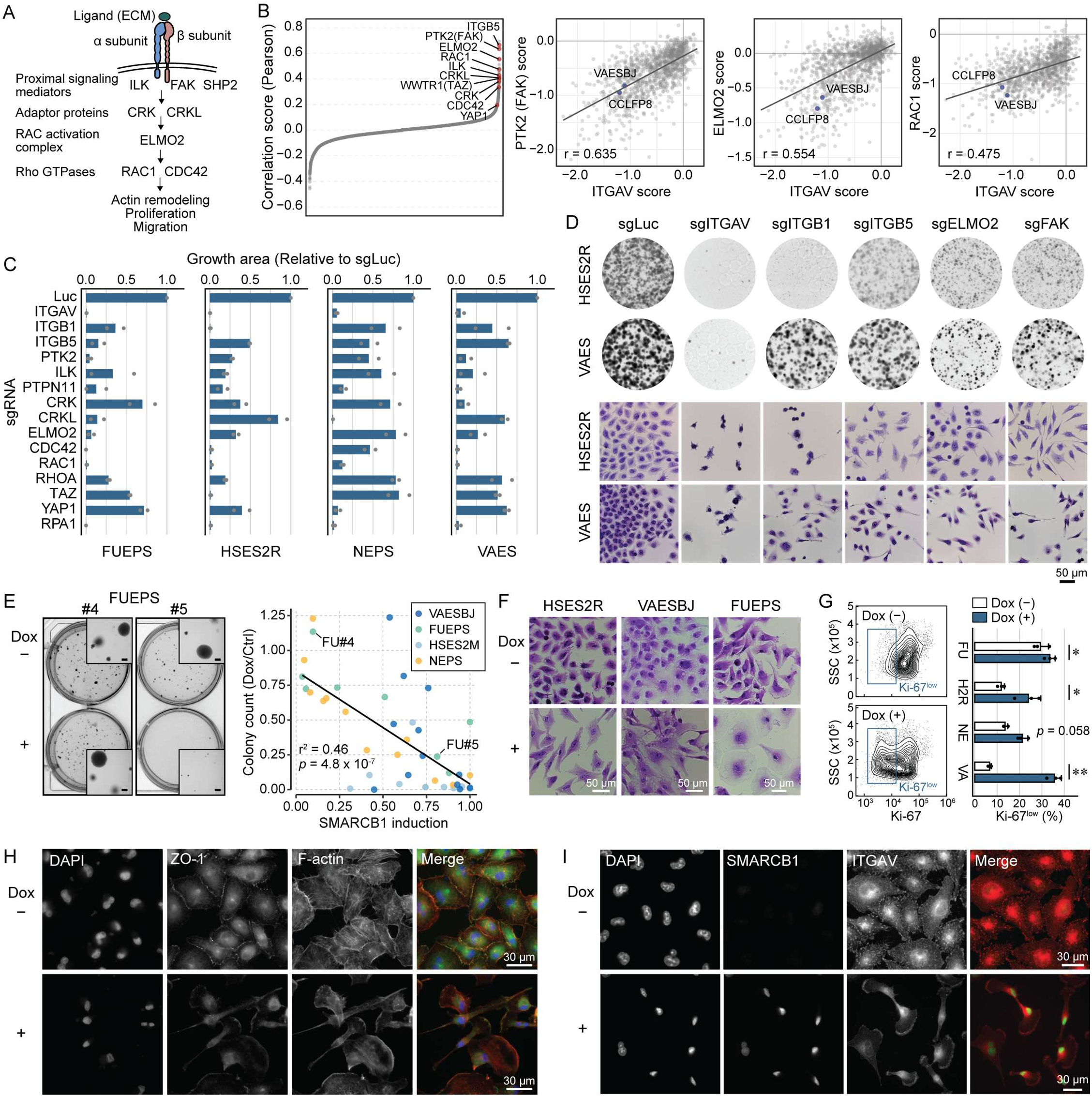
| Integrin signaling supports epithelioid proliferation in EPS. (A) Schematic of the integrin signaling cascade supporting proliferation in EPS. Signaling components identified as EPS dependencies are shown. ECM, extracellular matrix. (B) Correlation analysis of DepMap dependency scores for ITGAV. Highlighted genes are EPS dependencies identified in DepMap or our CRISPR screens. *r* indicates the Pearson correlation coefficient. (C) Effects of knockout of integrin signaling components on EPS cell growth. Growth area was quantified based on crystal violet staining. Plots show the mean of two independent experiments with individual replicates indicated. sgRNAs targeting firefly luciferase (Luc) and RPA1 were used as negative and positive controls, respectively. (D) Representative images showing impaired EPS growth and morphological changes after knockout of integrin signaling genes. (E) Soft agar assay in four EPS cell lines expressing Tet-ON SMARCB1. Left, representative growth phenotypes of two FUEPS clones with low (#4) or high (#5) doxycycline (Dox)-induced SMARCB1 expression. Bars in the inset images indicate 200 µm. Right, correlation between Dox-induced SMARCB1 expression and reduced colony-forming capacity. Values are from two independent replicates. (F) Morphological changes after SMARCB1 re-expression under standard adherent culture conditions. (G) Ki-67 expression in EPS cell lines without or with SMARCB1 restoration. The representative contour plots from VAES are shown. Data are mean ± SD. n = 3; \**p* < 0.05, \*\**p* < 0.01; paired two-tailed Student’s *t*-test. (H, I) Immunocytochemistry (ICC) showing the distribution of ZO-1, F-actin, and ITGAV in VAES cells without or with SMARCB1 re-expression.

To determine the link between SMARCB1 loss and integrin signaling dependence, we took a SMARCB1 add-back approach in EPS cells^18^. We employed a second-generation Tet-ON system to enable conditional SMARCB1 re-expression while minimizing supraphysiological expression (**Supplementary Fig. 3A, B**). For each of eight EPS lines (CCLFP8, ES2, FUEPS, HSES1, HSES2R, HSES2M, NEPS, VAES), we established 8–12 single-cell-derived subclones and assessed anchorage-independent growth in soft agar with or without doxycycline (Dox). In the four lines that formed macroscopically discernible colonies (FUEPS, HSES2R, NEPS, VAES), the degree of Dox-induced SMARCB1 expression inversely correlated with colony formation; highly induced clones showed marked loss of clonogenic potential (**Fig. 2E**, **Supplementary Fig. 3C, D**). SMARCB1 re-expression also reduced proliferation in adherent culture, with G1-phase accumulation and lower Ki-67 expression (**Fig. 2F, G**, **Supplementary Fig. 3E**), and shifted cells from compact clusters to a dispersed, more mesenchymal morphology resembling that observed after integrin perturbation **(Fig. 2F**, **Supplementary Fig. 3F**).

Restoration of SMARCB1 led to redistribution of tight junction proteins, including β-catenin, ZO-1, and claudins, which localized to cluster boundaries in EPS cells but became less prominent after restoration (**Fig. 2H**, **Supplementary Fig. 2E**). F-actin stress fibers changed from prominent bundles to a more diffuse cytoskeletal pattern. Notably, ITGAV localization shifted from a punctate distribution in clustered EPS cells to a polarized accumulation at lamellipodia-like structures (**Fig. 2I**, **Supplementary Fig. 2E**). Together, these findings suggest that loss of SMARCB1 promotes integrin signaling that supports clustered growth and proliferation in EPS.

### Loss of SMARCB1 and integrin signaling converge on activation of the oncogenic MYC program

To determine how integrin signaling supports EPS fitness, we first characterized transcriptional changes associated with SMARCB1 loss. For each EPS cell line (n=8), we analyzed Tet-ON SMARCB1 clones showing clear growth arrest or morphological changes after Dox-induced SMARCB1 re-expression (**Fig. 2E**, **Supplementary Fig. 3F**). Consistent with the role of the BAF complex in transcriptional activation, SMARCB1 re-expression broadly upregulated genes across all analyzed EPS lines and the RT line G402 (**Fig. 3A, B, Supplementary Fig. 4A, Supplementary Data 2**). The number of differentially expressed genes varied among lines, potentially reflecting differences in SMARCB1 induction and the threshold for restoring BAF activity. In contrast, SMARCB1 expression minimally affected the transcriptomes of RH4 rhabdomyosarcoma (RMS) and HT-1080 fibrosarcoma cells, supporting a specific perturbation of BAF function in EPS (**Supplementary Fig. 4A, Supplementary Data 2**).

**Fig. 3.**
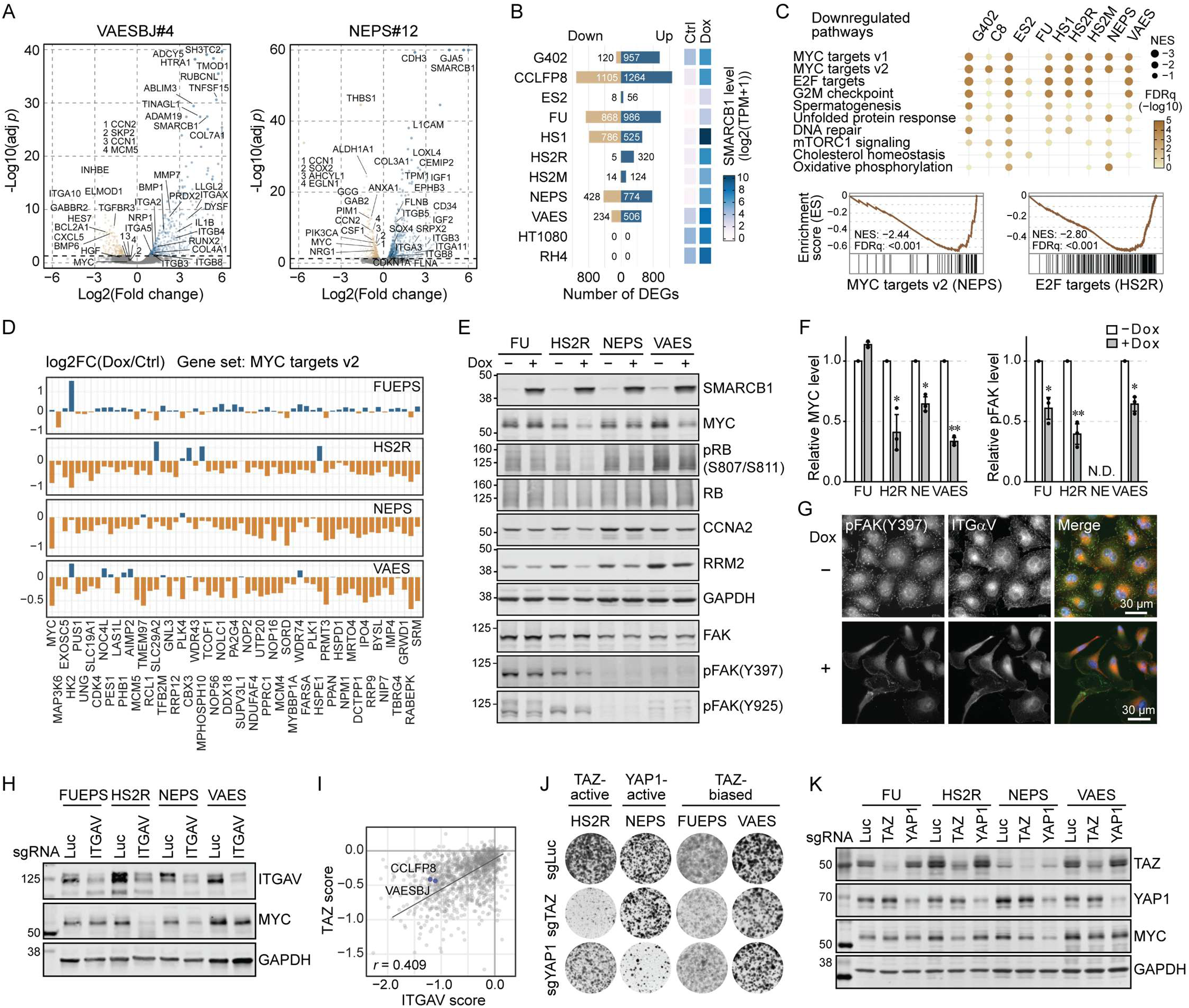
| Loss of SMARCB1 converges on dysregulation of integrin-MYC axis. (A) Transcriptomic changes after SMARCB1 restoration in EPS. Volcano plots of Tet-ON SMARCB1 clones from VAES and NEPS are shown. (B) Number of differentially expressed genes (DEGs) across eight EPS cell lines, the RT cell line G402, and two SMARCB1-proficient sarcoma cell lines (HT1080 and RH4). An adjusted *p*-value <0.05 was used as the cutoff to define DEGs. SMARCB1 expression levels are shown as log2(TPM+1). (C) Gene set enrichment analysis (GSEA) in G402 and eight EPS cell lines using Hallmark gene sets. Ten recurrently downregulated pathways and representative enrichment plots are shown. C8 = CCLFP8. (D) Expression changes in individual MYC target genes from the MSigDB MYC targets v2. Values are in log2 fold change (log2FC). (E) Western blot analysis of MYC, cell-cycle markers (RB, CCNA2, RRM2), and FAK. (F) Quantification of MYC and pFAK levels in EPS cells without or with SMARCB1 re-expression. n = 3; \**p* < 0.05, \*\**p* < 0.01; paired two-tailed Student’s *t*-test. N.D., not detected. (G) ICC showing reduced pFAK clusters in VAES cells that largely mirror the ITGAV distribution. (H) Western blot showing MYC downregulation after ITGAV knockout. (I) Positive correlation between TAZ and ITGAV dependency scores in DepMap. *r* indicates the Pearson correlation coefficient. (J) Representative images showing impaired EPS growth after TAZ or YAP1 knockout. (K) Western blot showing MYC downregulation after TAZ or YAP1 knockout in EPS cell lines.

Consistent with reduced proliferation after SMARCB1 re-expression, gene set enrichment analysis (GSEA) showed downregulation of E2F targets and G2M checkpoint genes (**Fig. 3C**). MYC targets, including MYC itself, were among the most strongly downregulated gene sets in seven of eight EPS lines, with FUEPS as the exception (**Fig. 3D**, **Supplementary Fig. 4B, C**). In support of these findings, western blotting after Dox treatment also showed reduced MYC and lower levels of cell-cycle progression markers, including phospho-RB (pRB), cyclin A2 (CCNA2), and ribonucleotide reductase 2 (RRM2) (**Fig. 3E, F**). These findings suggest that MYC-driven transcriptional programs are central to EPS fitness.

Ligand binding promotes integrin clustering and FAK autophosphorylation (pFAK), a key step in integrin outside-in signaling^19,20^. SMARCB1 re-expression decreased pFAK levels in EPS cells, although the magnitude varied among lines (**Fig. 3E, F**). In VAES cells, the pFAK signal was modest on western blotting but clearly detectable by immunocytochemistry and largely colocalized with ITGAV (**Fig. 3G**). After SMARCB1 re-expression, pFAK puncta became less prominent or redistributed to lamellipodia-like structures, consistent with altered ITGAV localization. Similar changes in pFAK distribution were observed in other EPS lines (**Supplementary Fig. 4D**). To test whether integrin signaling converges on MYC, we knocked out key pathway components. ITGAV KO reduced MYC in all EPS lines except FUEPS, matching the cell-line pattern observed after SMARCB1 re-expression (**Fig. 3H**). FAK, ITGB1, and ITGB5 KO produced similar effects (**Supplementary Fig. 4E**).

In DepMap, dependency on WWTR1 (TAZ) was highly correlated with dependency on ITGAV and FAK, suggesting functional coupling between integrin signaling and the Hippo pathway (**Fig. 2B**, **3I**, **Supplementary Fig. 4F**). Genetic knockout of YAP/TAZ strongly inhibited growth and reduced MYC expression in a mutually exclusive pattern across EPS lines, suggesting that YAP/TAZ may mediate integrin-driven MYC activation (**Fig. 3J, K**). Consistent with this model, canonical YAP/TAZ targets such as CCN1 (CYR61) and CCN2 (CTGF) were among the genes most strongly downregulated after SMARCB1 re-expression in NEPS cells (**Fig. 3A**, **Supplementary Fig. 4G**). Together, these findings support a model in which rewired integrin signaling converges on YAP/TAZ-MYC activation to promote EPS proliferation.

### Loss of SMARCB1 reshapes integrin receptor expression in EPS

To explore how SMARCB1 loss rewires integrin signaling, we analyzed genes upregulated after SMARCB1 restoration, which may include direct BAF targets. Although SMARCB1-responsive genes varied among cell lines, a recurrent set of genes was upregulated across all eight EPS lines (**Fig. 4A**). Pathway analysis showed strong enrichment of cell adhesion and extracellular matrix (ECM) remodeling (**Fig. 4B**). GSEA of individual lines also highlighted epithelial-to-mesenchymal transition (EMT), consistent with the mesenchymal shift after SMARCB1 re-expression (**Fig. 2F, 4C**). Upregulated genes included laminins (LAMB1, LAMB3), collagens (COL3A1, COL12A1), and ECM proteases (BMP1, HTRA1), providing a transcriptional basis for the morphological change. These findings suggest that loss of SMARCB1 promotes a shift to epithelioid phenotype in EPS.

**Fig. 4.**
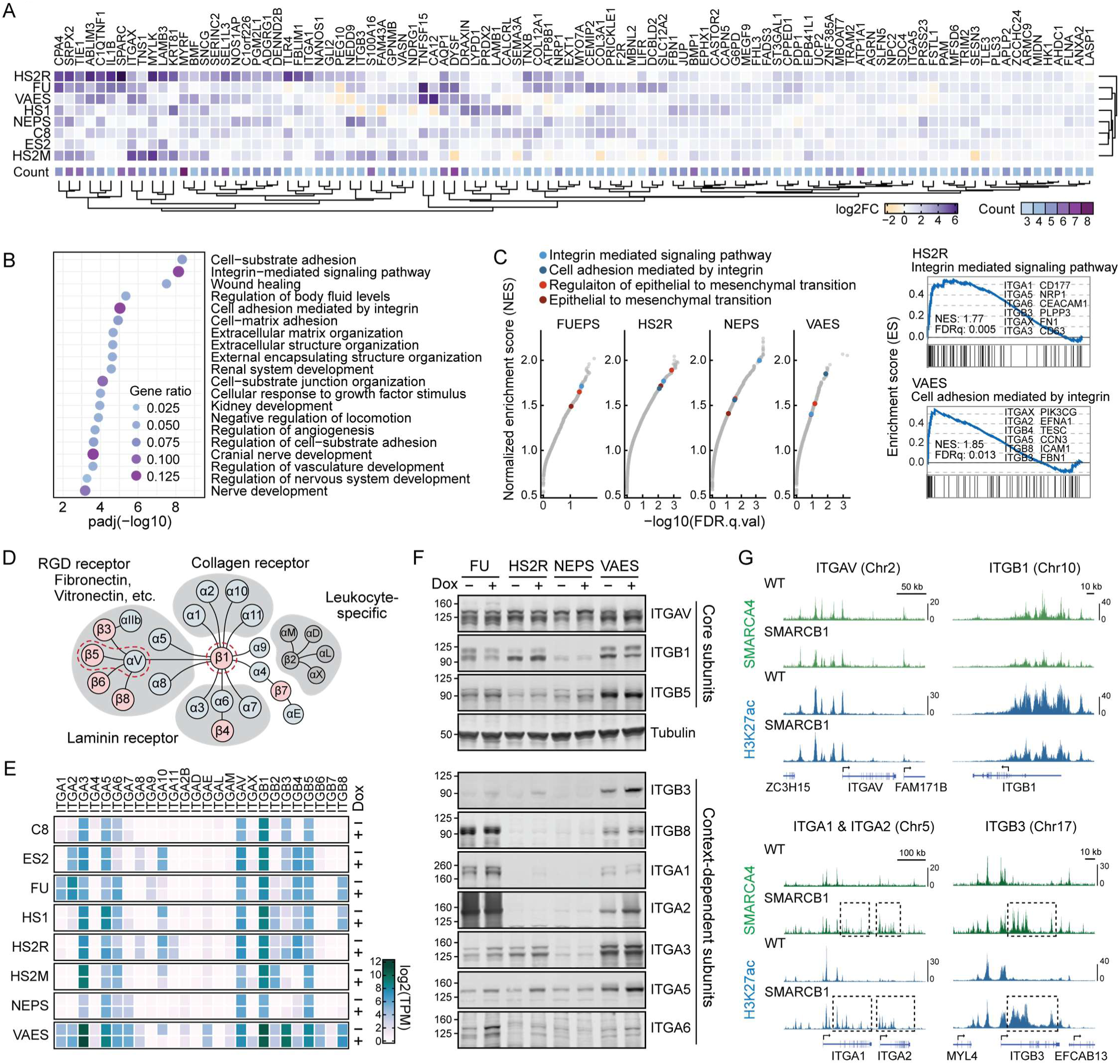
| Loss of SMARCB1 represses expression of context-dependent integrins while preserving core subunits. (A) Recurrently upregulated genes across eight EPS cell lines after SMARCB1 re-expression. Genes were ranked by DESeq2 *stat* score in each cell line, and recurrence scores were calculated as summed *stat* scores across lines. Fold changes for the top 100 genes are shown as a heatmap. Counts indicate the number of cell lines in which each gene was upregulated (Adjusted *p*-value <0.05). (B) Pathway enrichment of recurrently upregulated genes in EPS cell lines. The top 200 genes were analyzed using g:Profiler. (C) Pathways enrichment in each EPS cell line. GSEA was performed using MsigDB Gene Ontology Biological Process (C5) gene sets. Integrin signaling- and EMT-related pathways are highlighted; representative enrichment plots are shown on the right. (D) Integrin heterodimerization network. (E, F) mRNA- and protein-level expression of integrin receptor subunits. (G) ChIP-seq profiles of SMARCA4 and H3K27ac occupancy in VAES cells at integrin receptor gene loci (ITGA1, ITGA2, ITGAV, ITGB1, ITGB3). Dashed rectangles indicate gained peaks.

Notably, integrin signaling was also among the most significantly enriched pathways. Six receptor subunits (ITGA1, ITGA3, ITGA5, ITGA6, ITGAX, ITGB3) ranked among the top 200 recurrently upregulated genes, suggesting that specific integrin subunits are repressed in EPS (**Fig. 4A**, **Supplementary Fig. 5A**). This was unexpected given the strong integrin dependence in our CRISPR screens (**Fig. 1D, E**). We therefore generated a comprehensive expression profile of integrin receptor subunits in EPS. In the integrin heterodimerization network, ITGAV and ITGB1 are core nodes, with ITGB5 as the major ITGAV partner^21^ (**Fig. 2A and 4D**). ITGAV and ITGB5 were largely unchanged after SMARCB1 restoration, whereas ITGB1 was downregulated (**Fig. 4F**). In contrast, the restoration of SMARCB1 induced the expression of several context-dependent subunits, including ITGA1, ITGA2, ITGA3, ITGA5, ITGA6, ITGB3, and ITGB8.

To determine whether these integrin subunits are directly regulated by the BAF complex, we analyzed ChIP-seq data from VAES cells^10^. Consistent with the expression data, occupancy of SMARCA4, a component shared by the three major BAF complexes, was largely unchanged at the ITGAV locus regardless of SMARCB1 status but decreased at ITGB1 after SMARCB1 re-expression (**Fig. 4G**). H3K27ac closely mirrored SMARCA4 occupancy, linking BAF distribution to transcriptional activity. Conversely, SMARCB1 re-expression increased SMARCA4 recruitment and H3K27ac accumulation at upregulated genes, such as ITGA1, ITGA2, and ITGB3 (**Fig. 4G**, **Supplementary Fig. 5B**). Similar chromatin remodeling occurred at ECM-associated genes, including LAMB3, BMP1, and HTRA1. Together, these findings indicate that the loss of SMARCB1 reshapes the integrin receptor landscape by selectively repressing context-dependent subunits and ECM- associated genes while preserving the ITGAV-centered axis, potentially explaining the strong ITGAV dependency. Because restoration of SMARCB1 also induced regulators of integrin signaling, including NRP1, FLNA, and FLNB, additional mechanisms likely contribute to pathway rewiring^22,23^.

### CDKN2A (p16) is silenced genetically or epigenetically in EPS, potentially contributing to tazemetostat resistance

Targeting core integrin signaling nodes has been challenging because of toxicity in noncancer tissues^24,25^. To identify alternative therapeutic vulnerabilities, we revisited our druggable CRISPR screening data. Because none of the eight EPS cell lines exhibited a substantial growth disadvantage upon EZH2 KO (**Fig. 1D**), we tested the EZH2 inhibitor tazemetostat, and the recently developed EZH1/2 dual inhibitor valemetosat. The RT cell line G402 showed a moderate but discernible response to both compounds, whereas all EPS cell lines were generally resistant (**Fig. 5A**, **Supplementary Fig. 6A**). A similar contrast between RT and EPS models in response to tazemetostat has been reported by Kazansky et al.^26^, who observed frequent CDKN2A deletion and a lack of p16 induction following tazemetostat treatment in EPS cell lines. Consistent with these findings, p16 remained undetectable after tazemetostat treatment in all seven EPS lines tested, including five in which p16 levels had not previously been reported, despite a marked H3K27me3 reduction (**Fig. 5B**). Moreover, p16 KO was sufficient to confer resistance to EZH1/2 inhibition in G402, supporting a causal role for p16 in mediating drug resistance (**Fig. 5C**, **Supplementary Fig. 6B, C**).

**Fig. 5.**
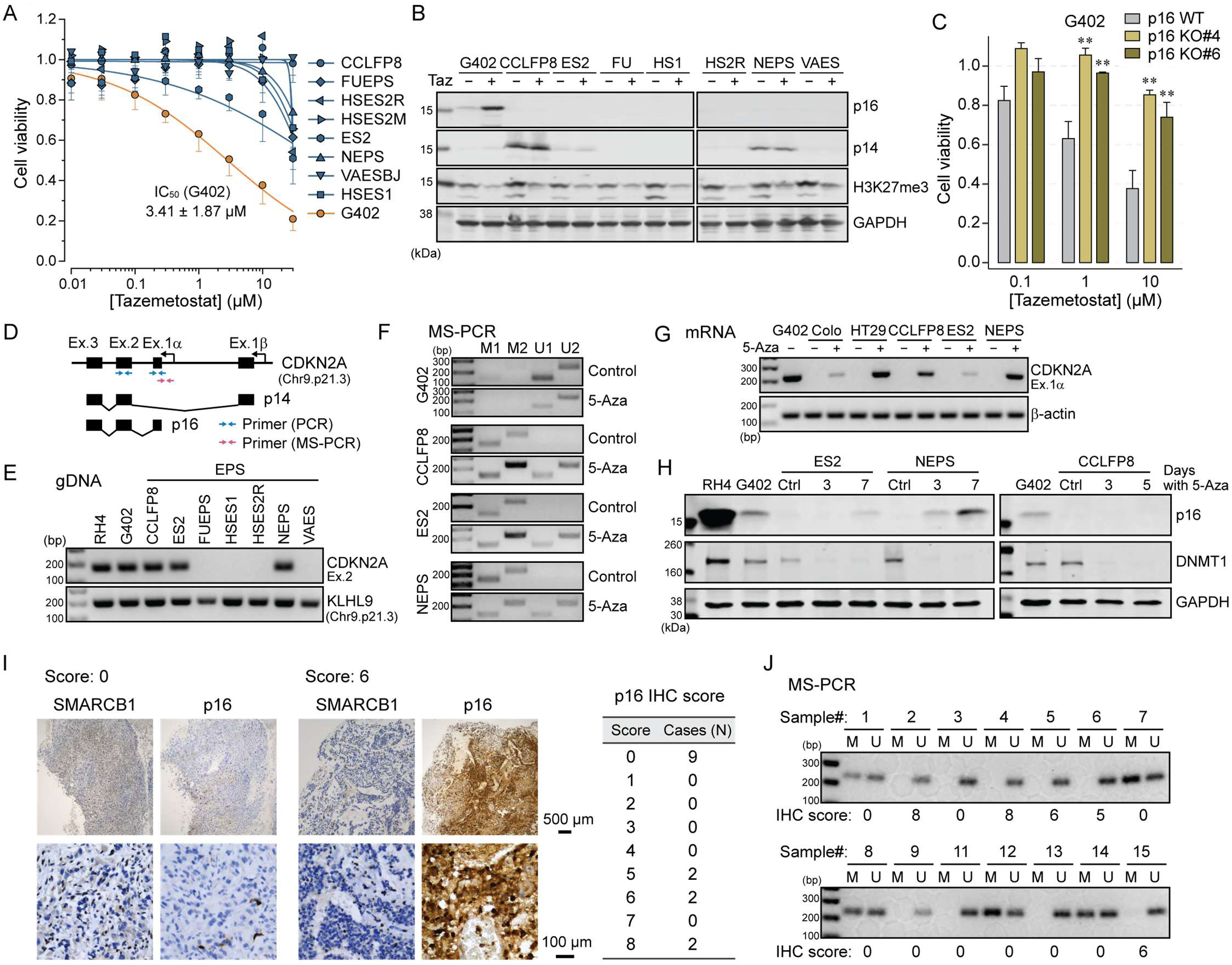
| p16 silencing and resistance to EZH1/2 inhibition in EPS. (A) Cell viability assay for tazemetostat (Taz). Eight EPS cell lines (blue) and the RT cell line G402 (orange) were treated with Taz for 7 days. (B) Protein expression of p16, p14 and H3K27me3 after 7-day treatment with Taz (5 µM). (C) Effect of p16 knockout on Taz-induced cytotoxicity in G402 cells. Two independent knockout clones were analyzed. \*\**p* <0.01. One-way ANOVA followed by Dunnett’s test. (D) Structure of CDKN2A. Transcript variants p14 and p16 are driven by distinct promoters. Horizontal arrows indicate primers used for genomic DNA PCR (gDNA-PCR), mRNA RT-PCR (E and G, blue), and methylation-specific PCR (MS- PCR) (F, red). (E) gDNA-PCR using primers targeting the shared exon (Ex. 2). KLHL9, a neighboring gene, served as a positive control. (F) MS-PCR in G402, CCLFP8, ES2 and NEPS cells. Bisulfite-treated gDNA was amplified using primers specific for methylated (M) and unmethylated (U) p16 promoter sequences. Two primer pairs (M1 and M2, or U1 and U2) were used for each methylation state. (G) Recovery of p16 transcript after 5-azacytidine (5-Aza) treatment. mRNA was analyzed by RT-PCR using primers for p16-specific exon 1α. Cells were treated with 3 µM 5-Aza for 7 days (all lines except CCLFP8) or 5 days (CCLFP8). Colon cancer cell lines COLO 205 (Colo) and HT-29 were used as methylation-positive controls. (H) Recovery of p16 protein with concomitant DNMT1 reduction after 5-Aza treatment. Treatment durations are as indicated. (I) Immunohistochemical (IHC) analysis of SMARCB1 and p16 in 15 EPS cases. Representative sections and summary of p16 scores are shown. (J) MS-PCR in primary EPS tumors. Samples from which gDNA were recovered (n = 14) were analyzed. Primer pairs M2 and U2 were used.

Clinical sequencing data for EPS have recently become available^14,27^ (**Supplementary Fig. 6D**). Despite frequent p16 loss in cell lines, CDKN2A deletion is relatively uncommon in primary EPS tumors (3-6%). Therefore, we investigated alternative mechanisms of p16 silencing that might contribute to the low response rate to tazemetostat in EPS patients^2^. Genomic PCR identified focal CDKN2A deletions in four EPS lines, whereas CCLFP8, ES2 and NEPS, as well as the positive controls RH4 and G402, retained an intact CDKN2A locus (**Fig. 5D, E**). Notably, p14, a CDKN2A transcript variant driven by a distinct promoter, was expressed in these three EPS lines, suggesting promoter-level p16 silencing (**Fig. 5B**). Methylation-specific PCR (MS-PCR) confirmed p16 promoter methylation in CCLFP8, ES2, and NEPS, and in two positive-control colon cancer cell lines^28^ (**Fig. 5F**, **Supplementary Fig. 6E**). Treatment with the DNA methyltransferase inhibitor 5-azacytidine (5- Aza) produced partial but discernible p16 promoter demethylation and mRNA recovery in all three EPS lines (**Fig. 5F, G**), and protein recovery, except in CCLFP8, for which five days was the longest tolerable treatment duration (**Fig. 5H**, **Supplementary Fig. 6F**).

Because inactivation of tumor suppressors may be overrepresented in cell lines through *in vitro* selection pressure, we next assessed p16 status in primary EPS samples. Immunohistochemistry (IHC) showed the absence of p16 staining in nine out of 15 cases, while the remaining six exhibited intermediate to high expression (Modified Allred score: 6–8) (**Fig. 5I, Supplementary Table 2**). MS-PCR in samples with successfully recovered gDNA (n = 14) confirmed p16 promoter methylation in five cases, all of which were negative for p16 staining in IHC (**Fig. 5J**). These findings suggest that inactivation of p16 might be more prevalent in EPS than previously recognized and may represent a key determinant of sensitivity to EZH2 inhibition.

### CDK4/6 are conserved dependencies in EPS

CRISPR screens revealed mutually exclusive CDK4 and CDK6 dependencies in EPS, a pattern that appears conserved across SMARCB1-deficient cancers (**Fig. 6A**). Considering the frequent p16 silencing and the clinical availability of CDK4/6 inhibitors, we investigated CDK4/6 as a therapeutic target in EPS. CRISPR competitive growth assays in NEPS and VAES showed strong depletion of sgRNAs targeting ITGAV, CDK2, CDK4, and CDK6, with the expected mutually exclusive dependency pattern between CDK4 and CDK6 (**Fig. 6B**). Consistent with the profile in DepMap, CDK6 depletion was also observed in the RMS cell line RH4, although to a lesser extent (**Supplementary Fig. 7A**). The dual CDK4/6 inhibitor palbociclib suppressed proliferation in all tested EPS lines (**Fig. 6C**, **Supplementary Fig. 7B**). Four lines (FUEPS, HSES2R, NEPS, VAES) had approximately 10-fold lower IC_50_ than the two RMS cell lines RD and RH4. Although CCLFP8, ES2, and HSES1 had IC_50_ values comparable to those of RMS models, they showed modest but significant growth suppression below 100 nM, with enlarged morphology suggestive of senescence (**Fig. 6C**, **Supplementary Fig. 7B**). Because ES2 and HSES1 proliferate slowly and CDK4/6 inhibition is cytostatic, short-term viability assays may underestimate drug response^29^ (**Supplementary Fig. 7C, D**). Accordingly, long-term clonogenic assays revealed marked sensitivity in CCLFP8, ES2, and HSES1, with substantial growth suppression in ES2 and HSES1 at palbociclib concentrations as low as 20 nM (**Fig. 6D**).

**Fig. 6.**
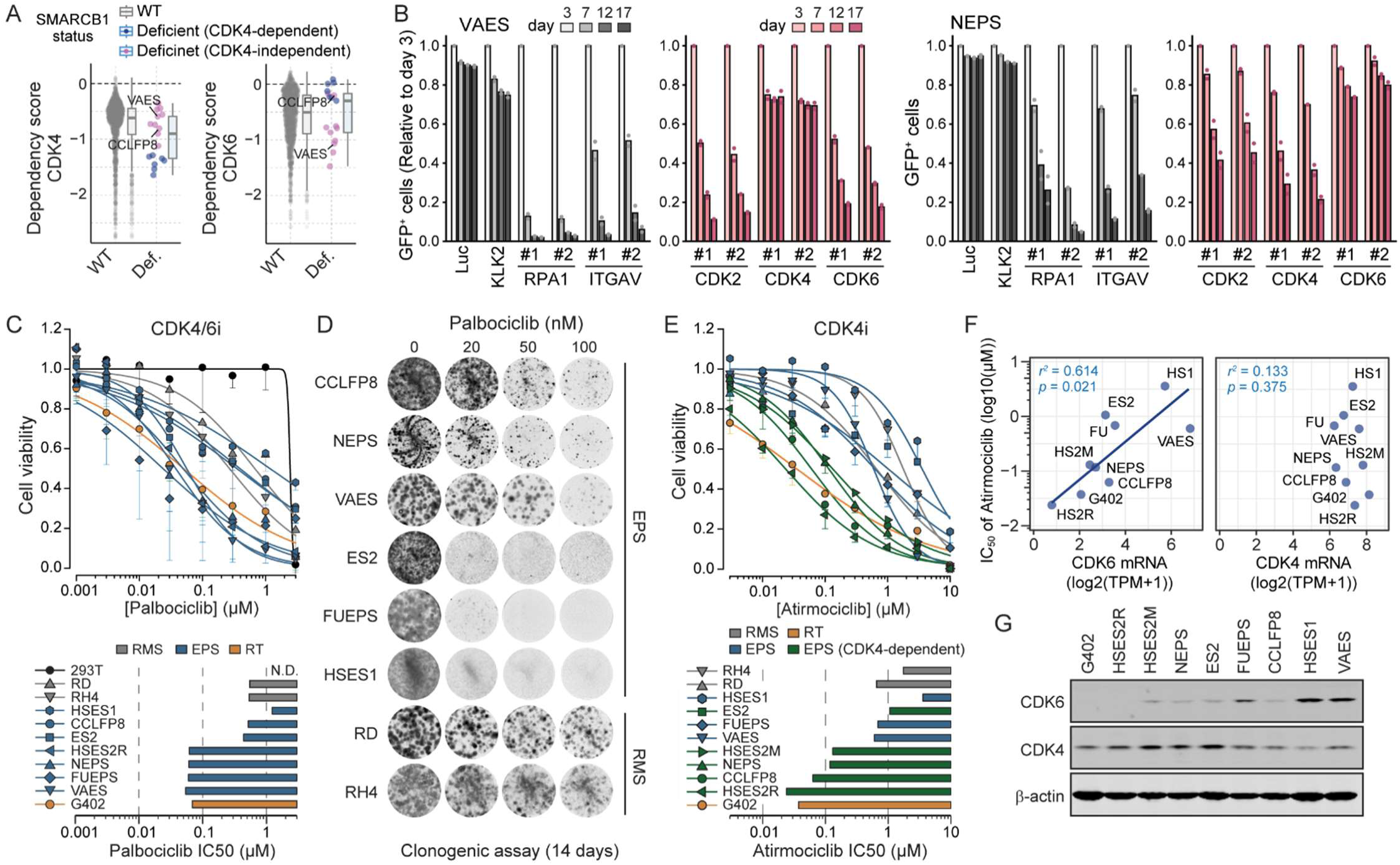
| CDK4 and CDK6 are targetable dependencies in EPS. (A) Two-class comparison of CDK4 and CDK6 dependency scores between SMARCB1 wild-type (n = 1,161) and SMARCB1-deficient (n = 17) cell lines. SMARCB1-deficient lines were further classified as CDK4-dependent or - independent to illustrate the mutually exclusive dependency pattern between CDK4 and CDK6. (B) CRISPR competitive growth assay in VAES and NEPS cells. sgRNAs were introduced into Cas9-expressing cells together with EGFP, and the EGFP-positive fraction was monitored by flow cytometry from day 3. Bars represent the mean of two independent experiments; dots indicate individual replicates. Luciferase (Luc) and the prostate-specific gene KLK2 were used as negative controls, and RPA1 as a lethal control. (C, E) Cell viability assay for (C) palbociclib and (E) atirmociclib. Cells were treated with each compound for 7 days. Bottom panels show IC_50_ values. In (E), EPS cell lines were classified as CDK4-dependent or -independent based on CRISPR screen data. HEK293T cells and cell lines from EPS (blue), RT (yellow), and rhabdomyosarcoma (RMS, grey) were used. n = 3, mean ± SD. (D) Clonogenic assay of long-term palbociclib treatment. Cells plated at clonogenic density were treated with palbociclib for 14 days and stained with crystal violet. (F) Correlation of atirmociclib sensitivity with CDK6, but not CDK4, mRNA levels in G402 and eight EPS cell lines. (G) Western blot analysis of CDK4 and CDK6 in G402 and eight EPS cell lines. CDK6 levels show greater cell line- dependent variability.

Atirmociclib is a recently developed selective CDK4 inhibitor with improved tolerability attributed to reduced hematopoietic toxicity^30^. EPS lines classified as CDK4-dependent in the CRISPR screens were markedly more sensitive to atirmociclib, with approximately 10-fold lower IC_50_ values compared with CDK4-independent EPS and RMS lines (**Fig. 6E**, **Supplementary Fig. 7E, F**). CDK6 expression negatively correlated with atirmociclib sensitivity, supporting CDK6 as a determinant of CDK4 dependency (**Fig. 6F, G**).

These findings prompted us to examine the expression of other CDK inhibitor proteins. p21 and p27 were modestly but consistently upregulated following SMARCB1 re-expression in EPS cells, which may reflect direct regulation by the BAF complex or an indirect post-translational mechanism mediated by SKP2 (**Supplementary Fig. 8A–D**).

Finally, we evaluated additional targets identified in the CRISPR screens. Drug response profiles were concordant with the screening results, with EPS cells showing sensitivity to CDK2 inhibition, whereas SMARCA4 and MDM2 inhibition had limited or genotype-dependent effects, respectively (**Supplementary Fig. 8E–H**). Resistance to MDM2 blockade was associated with oncogenic mutations in p53 in a subset of EPS cell lines, potentially explaining the reduced p21 levels (**Supplementary Fig. 8I**). Overall, these data reinforce loss of cell cycle control as a hallmark of EPS and support the evaluation of CDK-targeted strategies.

### *In vivo* efficacy of dual CDK4/6 inhibition and translational potential of CDK4-selective blockade

We next evaluated the therapeutic efficacy of CDK4/6 inhibition *in vivo*. The *in vivo* growth kinetics of most EPS cell lines have been poorly characterized, apart from the well-studied VAES. Therefore, we first tested their ability to form subcutaneous xenografts in NSG mice. We evaluated three lines with confirmed anchorage-independent growth in soft agar, as well as ES2, a PDX-derived cell line (**Supplementary Fig. 9A**). FUEPS, NEPS, and HSES2M formed tumors, with FUEPS and NEPS showing shorter latencies to palpable tumor formation. We then established VAES xenografts in athymic nude mice and NEPS xenografts in NSG mice to test palbociclib. Treatment with 100 and 150 mg/kg of palbociclib significantly delayed tumor growth in both models (**Fig. 7A**, **Supplementary Fig. 9B**). In support of on-target activity, palbociclib-treated tumors showed reduced pRB, Ki67, and CCNA2 (**Fig. 7B**).

**Fig. 7.**
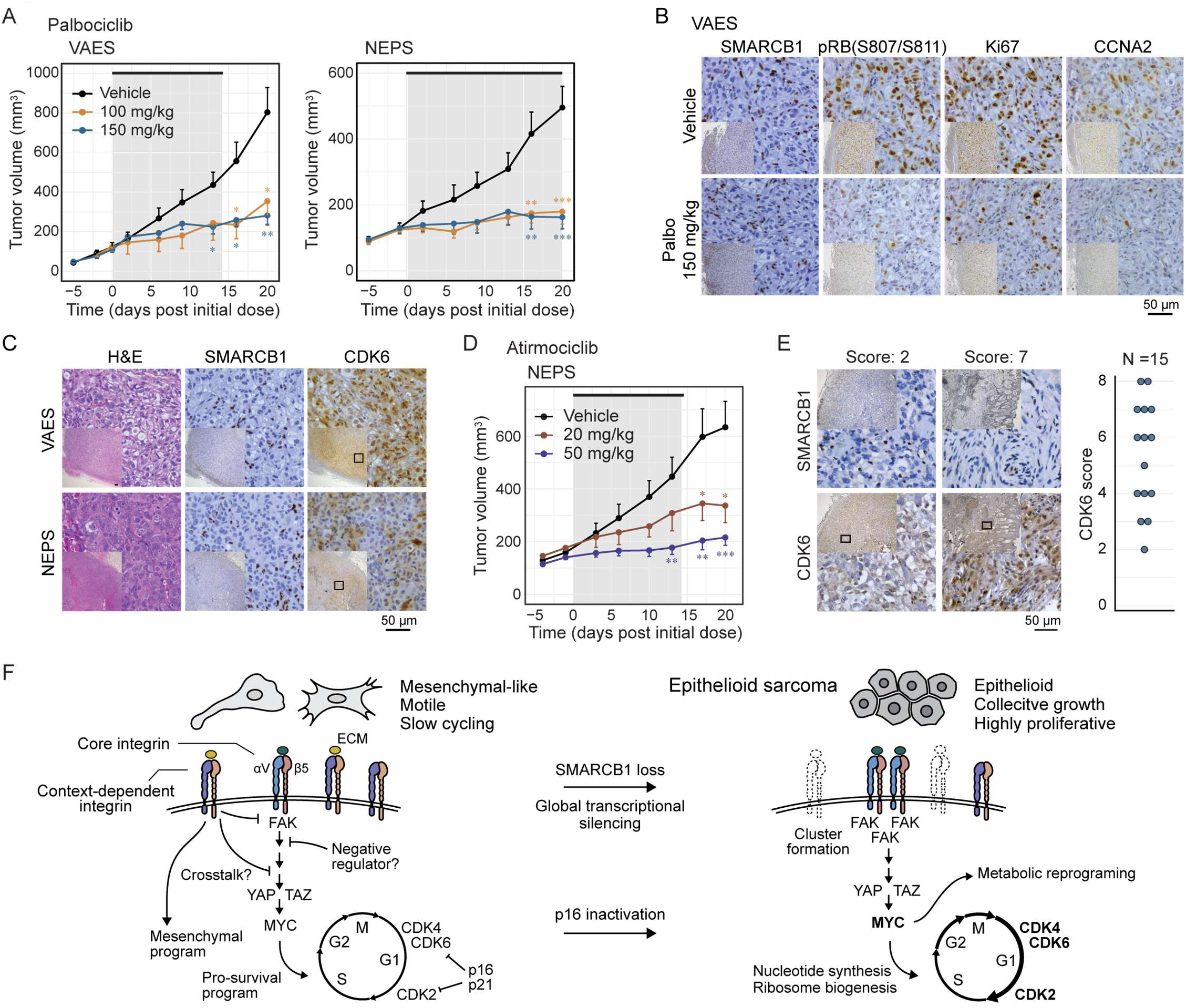
| In vivo efficacy of CDK4/6 blockade in EPS. (A) Tumor growth suppression by palbociclib in VAES-derived (left) and NEPS-derived (right) xenograft models. Cells were implanted subcutaneously in the right flank of athymic nude (VAES; n = 7-10) or NSG (NEPS; n = 8-10) mice. Vehicle or palbociclib (100 or 150 mg/kg) was administered daily by oral gavage for 14 days (VAES) or 20 days (NEPS). Values indicate mean ± SEM.\**p* < 0.05, \*\**p* < 0.01, \*\*\**p* < 0.001. One-way ANOVA followed by Dunnett’s test was used. (B) IHC showing reduced cell-cycle markers (pRB, CCNA2) and the proliferation marker (Ki-67) after palbociclib treatment. Representative images from one of three tumors are shown. SMARCB1 staining was used to identify tumor area. (C) CDK6 staining in VAES- and NEPS-derived xenografts. Sections were obtained from vehicle-treated mice in (A). (D) Efficacy of atirmociclib in NEPS-derived xenograft models (n = 8). Vehicle or atirmociclib (20 or 50 mg/kg) was administered twice daily for 14 days. Values indicate mean ± SEM. \**p* < 0.05, \*\**p* < 0.01, \*\*\**p* < 0.001. One-way ANOVA followed by Dunnett’s test was used. (E) CDK6 IHC in primary EPS tumors. Representative images of two scores are shown. (F) Proposed model of SMARCB1 loss-driven EPS development.

Differences in CDK6 expression were observed in VAES- and NEPS-derived tumors, with moderate CDK6 levels in NEPS tumors (**Fig. 7C**). Accordingly, atirmociclib produced a dose-dependent antitumor effect in NEPS xenografts, with a marked effect at 50 mg/kg, a clinically relevant dose^30^ (**Fig. 7D**, **Supplementary Fig. 9C, D**). Primary EPS tumors showed substantial variation in the expression of CDK6, with six of 15 cases showing weak-to-moderate nuclear staining (Modified Allred score ≤ 4) (**Fig. 7E**). These data demonstrate *in vivo* activity of CDK4/6 blockade in EPS and support a stratified treatment strategy in which CDK4-selective inhibition may improve tumor control in CDK6-low EPS.

## Discussion

In this study, through combined genetic screening and functional analyses in eight EPS cell lines, we identified oncogenic programs underlying the malignant phenotype of epithelioid sarcoma. Our findings support a model in which loss of SMARCB1 disrupts chromatin accessibility at integrin signaling genes, shifting the balance toward the oncogenic ITGAV-MYC axis (Fig. 7F). In parallel, p16 inactivation establishes a permissive cell-cycle state that sustains CDK4/6-dependent proliferation. These findings define EPS as a disease driven by coordinated dysregulation of integrin signaling and cell-cycle control (Fig. 7F).

In EPS, core integrin subunits including ITGAV and ITGB5 exhibited ubiquitous expression, whereas expression of other subunits was context-dependent. These variably expressed subunits were more susceptible to BAF regulation, contributing to SMARCB1-loss-driven perturbation of integrin receptor composition. ITGAV and its major partner ITGB5 remained largely unaffected by SMARCB1 status, whereas multiple alternative subunits were downregulated after SMARCB1 loss. Selective preservation of the ITGAV- centered axis may therefore explain the strong ITGAV dependency in our CRISPR screens. Crosstalk among integrin heterodimers, in which one subunit can interfere with another, could bias signaling toward specific downstream pathways in EPS^21^ (Fig. 7F). Loss of SMARCB1 may also alter signaling downstream of integrin receptors. NEPS illustrates this possibility, exhibiting limited changes in integrin receptor expression but pronounced deregulation of YAP1, a key downstream transcriptional regulator.

Our study suggests that inactivation of SMARCB1 in EPS is associated with a marked shift toward an epithelial-like state at both morphological and transcriptional levels, which may underlie the characteristic epithelioid immunophenotype of EPS. Because integrins are central to mechanotransduction and cytoskeletal organization, alterations in integrin repertoire may also contribute to this transition^20^. Additionally, loss of SMARCB1 downregulated genes associated with mesenchymal cell behavior, including ECM components and proteases. In sarcomas, which are generally considered mesenchymal tumors, the acquisition of epithelioid attributes has been linked to malignant progression through enhanced cell-cell cohesion, altered responses to extracellular cues, and paracrine activation of growth-promoting signaling pathways^31,32^. Epithelioid characteristics may therefore contribute to clonal expansion and tumor fitness rather than represent a passive feature of EPS. In parallel, an incomplete epithelial transition could result in relatively loose intercellular cohesion, potentially allowing cells to maintain some degree of collective behavior while remaining prone to shedding from the primary tumor and subsequent vascular dissemination. Retention of an intermediate epithelial-mesenchymal state, together with phenotypic plasticity, could provide a functional advantage during metastasis and contribute to the metastatic propensity of EPS.

Our study identifies deregulated cell-cycle progression as a shared hallmark of SMARCB1-deficient cancers, including both EPS and RT. However, the underlying mechanisms appear to differ. In RT, the BAF complex regulates CDK inhibitors such as p16 and p21; therefore, loss of SMARCB1 alone may be sufficient to drive cell-cycle progression^12,13^. In contrast, in EPS, the frequent occurrence of SMARCB1-independent p16 inactivation suggests that additional alterations might be required. Differences in cell of origin and lineage-specific BAF occupancy may contribute to these distinct outcomes. RT typically occurs in infants and children, whereas EPS predominantly affects adolescents and young adults and has a higher mutational burden^6^. These observations support a multistep model in which loss of SMARCB1 cooperates with secondary lesions, such as p16 inactivation, to drive EPS transformation. The need for additional oncogenic events may also help explain the later onset of EPS relative to RT. Thoracic SMARCA4-deficient undifferentiated tumors (SMARCA4- UTs) are BAF-perturbed cancers that occur in middle-aged adults^33,34^. Co-occurring driver alterations, including mutations in TP53, STK11, and KEAP1, have been documented in these tumors, providing an example consistent with this multistep model.

Although approximately 90% of EPS cases share SMARCB1 loss, substantial intertumoral heterogeneity with multiple histological subtypes is recognized. These include distal (classical) and proximal types, with the latter considered more aggressive^35,36^. Several other histological variants, such as fibrous and angiomatoid, have also been reported. The intrinsic heterogeneity may be reflected in our CRISPR screens, which showed marked cell line-specific dependency profiles. Cell of origin and cooperating oncogenic alterations may jointly shape this variation. Cell line-specific vulnerabilities included highly tractable targets such as MET and FGFR1. These findings suggest that integrin signaling and CDK4/6 dependency may represent common EPS vulnerabilities, while lineage- or genotype-specific dependencies offer additional opportunities for personalized treatment. An important consideration is that six cell lines used in this study were established from recurrent or metastatic tumors. Thus, we cannot rule out the possibility that some of the findings in this study reflect features of advanced disease. Validation in a broader set of cell lines and additional models will be important to substantiate the proposed concept and facilitate prioritization of therapeutic targets.

We observed mutually exclusive dependencies on CDK4 and CDK6, with CDK6 expression emerging as a potential biomarker of CDK4-dependent EPS. Dual CDK4/6 inhibitors such as palbociclib benefit some patients with breast cancer, but hematologic toxicity associated with CDK6 inhibition can limit dosing and sustained tumor control. The CDK4-selective inhibitor atirmociclib has shown improved tolerability and antitumor efficacy in preclinical breast-cancer models and is currently under evaluation in Phase III clinical trials^30^. Our findings suggest that CDK4-selective inhibition could enable deeper, more durable responses in CDK6-low EPS while reducing toxicity associated with dual CDK4/6 blockade.

In conclusion, systematic functional interrogation of EPS identified oncogenic programs central to its biology. SMARCB1 inactivation rewires integrin signaling, biasing activation toward an integrin-MYC axis that, together with deregulated cell-cycle progression, shapes the malignant state. Further studies are required to define how integrin crosstalk and associated proteins contribute to this oncogenic rewiring. Optimization of CDK4/6- targeted therapies and development of subtype-specific treatment strategies are also important directions for future research. Overall, this study provides mechanistic insight into the molecular basis of EPS and establishes a framework for rational development of targeted therapeutic strategies for this rare but aggressive sarcoma.

## Methods

### Cell culture

CCLFPEDS0008T (CCLFP8, HCM-BROD-0053-C49), COLO 205, G402, HT-1080, HT-29 and VAESBJ were purchased from American Type Culture Collection (ATCC). HSES1, HSES2R and HSES2M were obtained from RIKEN BioResource Research Center. FUEPS was kindly provided by Dr. Hiroshi Iwasaki^37^. NEPS was kindly provided by Dr. Hiroyuki Kawashima^38^. ES2 was established as described previously^39^. Human embryonic kidney (HEK) 293T cells were purchased from Takara Bio. RD and RH4 were kindly provided by Dr. Corinne Linardic^40^. All cell lines, except CCLFP8, were maintained in Dulbecco’s Modified Eagle Medium (DMEM) supplemented with 5% fetal bovine serum (FBS) and 100 U/mL penicillin-streptomycin (all from Thermo Fisher Scientific). CCLFP8 was maintained in Skeletal Muscle Cell Growth Medium-2 (SkGM-2) supplemented with SkGM BulletKit (both from Lonza). Cells were cultured at 37°C in a humidified 5% CO_2_ atmosphere and routinely tested for mycoplasma contamination using the MycoAlert Detection Kit (Lonza).

### Compounds

Palbociclib isethionate (S1579) and tazemetostat (S7128) were purchased from Selleck Chemicals. Atirmociclib (HY-139450), AU-15330 (HY-145388), idasanutlin (HY-15676), INX-315 (HY-162001), and valemetostat tosylate (HY-109108A) were purchased from MedChemExpress. Doxycycline hyclate was purchased from MilliporeSigma. All selection antibiotics were purchased from InvivoGen.

### Antibodies

Antibodies used in this study are listed in **Supplementary Table 3**.

### Plasmids

To construct pLV-Hygro-2A-mCherryNLS, the tandem sequence encoding the hygromycin resistance gene and mCherry linked by a P2A peptide was PCR-amplified from pCF525-EF1a-Hygro-P2A-mCherry-lenti (Addgene #115796, gift from Jennifer Doudna) and cloned into the pLV-EF1-IRES-Neo vector (Addgene #85139, gift from Tobias Meyer). The mCherry sequence was then replaced with a mCherry sequence flanked by the MYC nuclear localization signal (NLS). lentiCas9-Hygro was constructed by replacing the blasticidin resistance cassette in lentiCas9-Blast (Addgene #52962, gift from Feng Zhang) with a hygromycin resistance cassette.

For the sgRNA expression vector, sgRNA target sequences were cloned into lentiGuide-Puro (Addgene #52963, gift from Feng Zhang) by oligonucleotide annealing and ligation. All cloning enzymes were purchased from New England Biolabs (NEB). sgRNA target sequences used in this study are listed in **Supplementary Table 4**.

### Generation of Tet-ON SMARCB1 cell lines

EPS cell lines with doxycycline-inducible SMARCB1 expression were generated using pLXI_TRC401 SMARCB1 (Addgene #111182, gift from William Hahn), which carries the Tet-Advanced second-generation Tet- ON system. The Tet3G system (Takara Bio, #631187) was also used to express SMARCB1 in RH4 and HT- 1080 cells. The SMARCB1 coding sequence was PCR-amplified from pLXI401 and cloned into pLVX-TRE3G to generate pLVX-TRE3G-SMARCB1. Cells were sequentially transduced with pLVX-Tet3G and pLVX-TRE3G- SMARCB1.

### Lentiviral infection

Lentiviral vectors were produced in HEK293T cells. Transfer vectors were co-transfected with the packaging plasmids pCMV-dR8.2 dvpr (Addgene #8455) and pCMV-VSV-G (Addgene #8454, both gifts from Bob Weinberg) at a 3:2:1 weight ratio using Lipofectamine 2000 (Thermo Fisher Scientific). Medium was replaced 12-24 h after transfection, and supernatant was collected 36-48 h after transfection. For sgRNA library production, viral supernatant was collected twice on consecutive days.

### DepMap data analysis

DepMap data^41^ (version 25Q2) were used to analyze and visualize genome-wide CRISPR screen data. Two-class comparisons of SMARCB1 wild-type and SMARCB1-deficient cell lines were performed using the Custom Analyses tool on the DepMap website. Cell lines annotated as Embryonal Tumor, Rhabdoid Cancer, or Epithelioid Sarcoma in OncotreePrimaryDisease were considered SMARCB1-deficient for this analysis. To identify dependencies selective for the EPS cell lines CCLFP8 and VAESBJ, we selected genes with dependency scores at least 0.2 below the pan-cancer average.

### Druggable gene-targeted CRISPR screen

Druggable genes were selected from the Pharos platform^42^. Genes classified as Tclin or Tchem were included, yielding 2600 genes (**Supplementary Table 2**). Genes encoding BAF and ISWI complex components (n = 26) and lethal controls (n = 30) were also included. sgRNAs were designed according to Joung et al.^43^, with six sgRNAs per gene. A pooled oligonucleotide library was synthesized by Twist Bioscience and cloned into lentiGuide-Puro by Gibson Assembly (NEB). Cas9 was introduced into EPS cell lines using lentiCas9-Blast, followed by single-cell cloning. To assess Cas9 activity in bulk populations and single-cell clones, cells were transduced with an sgRNA targeting the AAVS1 locus and selected with puromycin. Genomic DNA (gDNA) was extracted 7-10 days after transduction using the DNeasy Blood and Tissue Kit (Qiagen). An approximately 500 bp fragment spanning the sgRNA target site was PCR-amplified and analyzed by Sanger sequencing. Editing efficiency and indel patterns were inferred using the ICE tool (Synthego).

To initiate the screen, a predetermined volume of the sgRNA library was added to culture medium containing 4 µg/mL polybrene (MilliporeSigma) to achieve a 30-40% infection rate. Puromycin selection began 32-36 h after infection. On day 4, cells were harvested; a portion was retained for gDNA extraction to determine reference sgRNA representation (P0) and the remainder was reseeded at ∼20% confluence. Cells were passaged four additional times to achieve 10-12 doublings and then harvested for gDNA extraction to determine terminal sgRNA representation (P5). Screens lasted 17-20 days for NEPS, HSES2R, and HSES2M and 25-30 days for ES2, HSES1, and FUEPS. Coverage of 400× was maintained throughout. After gDNA extraction, NGS libraries were prepared by a two-step PCR method^44^ and sequenced to a depth corresponding to at least 1,000x coverage of total sgRNAs.

### Analysis of CRISPR screen

sgRNA counts were obtained using a Python script from Joung et al.^43^. Subsequent bioinformatic analyses and visualization were performed in R. For log transformation, a sample-specific pseudocount equal to the median sgRNA count divided by 20 was added to each sgRNA count. Counts were normalized by sequencing depth, and P5/P0 ratios were calculated and log_2_-transformed. Gene-level depletion scores were calculated as the mean log_2_ fold change of the six sgRNAs targeting each gene. Scores were calibrated against 260 essential and 200 nonessential genes to approximate the DepMap scale and facilitate identification of selective rather than core cancer dependencies. Median scores remained ∼0.2 lower than that in DepMap; therefore, selective dependencies were defined as scores at least 0.4 below the DepMap average rather than the 0.2 threshold used for VAES and CCLFP8. Functional enrichment analysis was performed using the STRING database^45^.

The 130 genes identified as dependencies in at least three cell lines were analyzed with default settings except that the minimum required interaction score was set to the highest confidence level (0.9). The resulting network data was visualized using Cytoscape^46^.

### CRISPR competitive growth assay

sgRNA target sequences were cloned into the LRG sgRNA expression vector carrying EGFP as a reporter (Lenti_sgRNA_EFS_GFP, Addgene #65656, gift from Christopher Vakoc). Cas9-expressing cells were transduced with the LRG vector, and the percentage of GFP-positive cells was measured using a CytoFLEX Flow Cytometer (Beckman Coulter) 3, 7, 12, and 17 days post-infection. To ensure high Cas9 activity in the polyclonal population, Cas9 transduction was repeated with lentiCas9-Blast and lentiCas9-Hygro. sgRNAs targeting firefly luciferase and the prostate-specific gene KLK2 were used as negative controls, and an sgRNA targeting RPA1 was used as a lethal control.

### Cell viability assay

To minimize the effect of contact inhibition, plating density was optimized for each cell line so that cells reached 90-100% confluence on the day of quantification. Growth rate and cell size were the two major determinants for the optimized plating density. Cells were seeded in 96-well plates at 500 cells/well (VAESBJ, 293T), 1,000 cells/well (G402, HSES2R, HSES2M, NEPS, RD, RH4) and 2,000 cells/well (CCLFPEDS0008T, ES2, FUEPS), or 4,000 cells/well (HSES1). Unless otherwise indicated, cells were treated with test compounds for 7 days and cell viability was measured using CellTiter-Glo (Promega) according to the manufacturer’s protocol. On day 4, half of the culture medium was replaced with fresh medium containing the indicated compound concentrations. Dose-response curves were fitted in OriginLab using a logistic function with upper and lower values fixed at 1 and 0, respectively. IC_50_ values were calculated for each biological replicate and averaged across at least three replicates per compound.

### Clonogenic assay

Cells were seeded in 6-well plates at 500 cells/well (VAESBJ, 293T), 1,000 cells/well (G402, HSES2R, HSES2M, NEPS, RD, RH4), 2,000 cells/well (CCLFPEDS0008T, ES2, FUEPS), or 4,000 cells/well (HSES1) and cultured for 2 weeks. Cells were stained with 0.01% crystal violet (MilliporeSigma) in 10% ethanol. Plates were scanned using a ChemiDoc imaging system (Bio-Rad), and cell-covered area was quantified in ImageJ. For clonogenic assays after CRISPR-mediated knockout, cells were transduced with sgRNA-expressing lentivirus and selected with puromycin for 2 days beginning 1 day after transduction. Selected cells were then collected and seeded for clonogenic assays.

### Growth kinetics analysis

Growth kinetics of cell lines were measured using the Incucyte S3 live-cell imaging system (Sartorius). Cells stably expressing nuclear-localizing mCherry were generated using pLV-mCherryNLS-hygro. Cells were seeded at 2-5 × 10^4^ cells/well in 24-well plates, and brightfield and red-fluorescence images were acquired every 12 h. For curve fitting, data points between 500 and 6,000 cells, plus one point outside each boundary, were fitted to the Gompertz function (SGompertz) in OriginLab. The time required for cell numbers to increase from 500 to 4,000, corresponding to three population doublings (3 × T2), was calculated using the uniroot function in R.

### Soft agar colony formation assay

Cells were cultured in a soft agar medium consisting of 0.4% and 0.7% agarose in the top and bottom layers, respectively. Briefly, low-melt agarose (Lonza) was dissolved in water at 2.4% and autoclaved. On the day of plating, the agarose was remelted and maintained at 45°C. For the bottom layer, agarose was diluted to 0.7% with prewarmed culture medium and dispensed into 6-well plates (1.5 mL/well). Cells were suspended in culture medium at 6,000 cells/mL. To prepare the top layer containing cells, a 1.2% agarose solution was prepared and maintained at 45°C. Equal volumes (1 mL each) of 1.2% agarose, culture medium, and cell suspension were mixed in this order to minimize heat shock and overlaid onto the solidified bottom layer at 1.5 mL/well, yielding 2,000 cells/well. After the agarose had solidified, 300-400 µL culture medium was added per well. For doxycycline (Dox) treatment, 2 µg/mL Dox was added to both layers. Cells were cultured for 4-6 weeks, with the overlying medium replaced weekly.

### Western blotting

Cells were collected by trypsinization followed by neutralization with complete culture medium. After washing with PBS, cells were lysed in RIPA buffer containing cOmplete protease inhibitor cocktail (Roche). PhosSTOP phosphatase inhibitor cocktail (Roche) was added when phosphorylated proteins were analyzed. After sonication and centrifugation, supernatants were collected and protein concentrations were determined using the Pierce BCA Protein Assay Kit (Thermo Fisher Scientific). Unless otherwise noted, lysates were diluted in RIPA buffer and mixed with 4× sample buffer (LI-COR) to a final protein concentration of 1-2 µg/uL. Typically, 10-20 µg protein was separated on precast SDS-PAGE gels (Bio-Rad or Thermo Fisher Scientific) at 150 V for 50-60 min and wet-transferred to nitrocellulose membranes. Standard transfer conditions were 100 V for 45 min; proteins ≤ 30 kDa were transferred at 80 V for 30 min. Membranes were blocked with Intercept Blocking Buffer (LI-COR) for 30 min at room temperature. Primary and fluorescent dye-conjugated secondary antibodies (LI-COR) were diluted in a 1:1 mixture of TBST and Intercept Blocking Buffer. Membranes were incubated with primary antibodies overnight at 4°C and secondary antibodies for 30 min at room temperature. Membranes were scanned using an Odyssey Imager (LI-COR). Band intensities were quantified using ImageJ.

### Animal experiments

Animal experiments were performed according to the guidelines of the Animal Resource Center (ARC) at the University Health Network (UHN), under the approved protocol (#6825). Male and female athymic nude mice (CrTac:NCr-*Foxn1^nu^*, Taconic Biosciences) and NSG mice (NOD.Cg-*Prkdc^scid^ Il2rg^tm1Wjl^/SzJ*, The Jackson Laboratory) were used. Five million cells were suspended in a 1:1 mixture of PBS and Matrigel (Corning) and injected subcutaneously into the right flank of athymic nude mice (VAES) or NSG mice (all other cell lines). Tumor length (*L*) and width (*W*) were measured twice weekly using a digital caliper, and tumor volume was calculated as *L* x *W*^2^/2. When mean tumor volume reached 75-100 mm^3^ for VAES or 125-175 mm^3^ for NEPS, mice were randomized into three groups. A stock solution of palbociclib isethionate (Selleck) was prepared in distilled water and diluted in 50 mM NaCl before use; palbociclib was administered once daily by oral gavage. A stock solution of atirmociclib (MedChemExpress) was prepared in DMSO and suspended in 0.5% methylcellulose containing 0.1% Tween 80 before use; atirmociclib was administered twice daily by oral gavage. Experimental endpoints were a tumor length > 1.5 cm or a tumor volume > 1000 mm^3^. Humane endpoints included tumor ulceration, severe body-weight loss (> 20%), or signs of poor general condition, including hunched posture, reduced activity, or abnormal breathing. Treatment continued for 14 or 20 days as indicated in the figure legends, and mice were monitored for 50 days after treatment initiation. Tumors reaching an endpoint were collected and fixed in 10% formaldehyde overnight for immunohistochemistry. If tissue collection occurred after the treatment period, mice were re-treated with the corresponding drug for two consecutive days before tissue collection.

### Primary EPS samples

Fifteen formalin-fixed paraffin-embedded specimens archived at Mount Sinai Hospital (Toronto) were used for IHC analysis under a Research Ethics Board (REB)-approved protocol (study ID 17-0103-E). All 15 samples had a confirmed diagnosis of EPS based on examination of hematoxylin and eosin (H&E)-stained sections. All were confirmed to be SMARCB1-negative by IHC. Patient information is provided in Supplementary Table 2.

### Immunohistochemistry (IHC)

Formalin-fixed tissues were paraffin-embedded and sectioned at 4 µm. After deparaffinization and rehydration, sections were treated with 3% hydrogen peroxide for 15 min at room temperature. Antigen retrieval was performed in citric acid-based retrieval solution (pH 6.0, Vector Laboratories) by microwave heating for 2 min followed by incubation at 95-100°C in a rice cooker for 30 min. Sections were blocked with 10% normal goat serum in PBS containing 0.1% IGEPAL CA-630 (PBST) for 30 min at room temperature and incubated overnight at 4°C with primary antibodies diluted in PBST containing 2% normal goat serum. Sections were then incubated with biotinylated goat secondary antibodies in the same diluent for 30 min, followed by ABC reagent (Vector Laboratories) for 30 min at room temperature. Signals were developed using a DAB Staining Kit (Vector Laboratories). Immunohistochemical staining was scored by the Allred method^47^.

### Immunocytochemistry (ICC)

Cells were seeded onto fibronectin-coated coverslips and were cultured for 5 days without or with 2 µg/mL Dox. Cells were fixed in methanol for 10 min at 4°C. After blocking with PBST containing 2% bovine serum albumin (BSA), cells were incubated with primary antibodies overnight at 4°C and fluorescent dye-conjugated secondary antibodies for 30 min at room temperature. Primary and secondary antibodies were diluted in PBST containing 0.2% BSA. Images were acquired using a Zeiss Axio Observer microscope (Zeiss).

### PCR

PCR was performed on gDNA, complementary DNA (cDNA), or bisulfite-treated gDNA as follows. gDNA was extracted using the DNeasy Blood & Tissue Kit. PCR was performed using GoTaq G2 Master Mix (Promega) with 20 ng template gDNA in a 20-30 µL reaction volume. Total RNA was extracted using the RNeasy Plus Kit (Qiagen) according to the manufacturer’s protocol, with on-column DNA digestion using the RNase-Free DNase Set (Qiagen). Reverse transcription was performed using the iScript cDNA Synthesis Kit (Bio-Rad). cDNA corresponding to 50 ng of the original RNA was amplified using GoTaq G2 Master Mix. For methylation-specific PCR (MS-PCR), DNA was bisulfite-treated using the EZ DNA Methylation Kit (Zymo Research) and amplified by hot-start PCR using ZymoTaq PreMix (Zymo Research). Primers for methylated and unmethylated p16 promoters were from Herman et al^48^. Primer sequences and annealing temperatures are listed in **Supplementary Table 5**. Tumor-tissue gDNA was extracted using the Quick-DNA/RNA FFPE Miniprep Kit (Zymo Research).

### Transcriptome analysis

RNA was extracted as described above. After ribosomal RNA depletion using the QIAseq FastSelect rRNA HMR kit (Qiagen), sequencing libraries were prepared using the NEBNext Ultra II RNA Library Preparation Kit for Illumina (NEB) and sequenced on an Illumina NovaSeq platform in a 2 × 150 bp configuration to a target depth of 20-40 million reads per sample. Raw FASTQ files were processed with fastp^49^ for adapter trimming and quality control, then aligned and quantified using STAR^50^ and RSEM^51^ with default parameters. Differential expression analysis was performed using DESeq2; transcripts with an average count < 5 across samples were excluded. For pathway enrichment analysis, genes were ranked by DESeq2 *stat* scores and analyzed by Gene Set Enrichment Analysis (GSEA) using the MSigDB Hallmark (H) and Gene Ontology Biological Process (C5) gene sets with a gene-set size cutoff of 50-500 genes. To identify recurrently upregulated genes following SMARCB1 re-expression, genes were ranked by summed *stat* scores across EPS cell lines and the top 200 genes were analyzed for pathway enrichment using g:Profiler with a gene-set cutoff of 50-500 genes^52^.

### Generation of p16 knockout clones

To generate CDKN2A (p16) knockout clones while minimizing off-target editing, we used Cas9 nickase (Cas9n). A pair of sgRNAs targeting exon 1α of CDKN2A was designed using CHOPCHOP^53^ and sequentially cloned into AIO-puro (Addgene #74630, gift from Steve Jackson) using the BbsI and BsaI restriction enzymes. The resulting plasmid was transfected into G402 cells using Lipofectamine 2000. One day after transfection, cells were selected with puromycin for 3 days, expanded, and seeded in 96-well plates by limiting dilution to obtain single-cell-derived clones. sgRNA target sequences are listed in Supplementary Table 1. Candidate clones were validated by western blotting.

### Cell cycle analysis

Cells were fixed in 70% ethanol for 15 min at room temperature and were incubated with a propidium iodide (PI) solution (Thermo Fisher Scientific) for 20 min at room temperature. The proportions of cells in G1, S, and G2/M phases were quantified using FlowJo (BD Biosciences). For Ki-67 staining, fixed cells were incubated with FITC-conjugated anti-Ki-67 antibody diluted in PBS containing 0.3% BSA for 30 min at room temperature. All flow cytometric analyses were performed using a CytoFLEX Flow Cytometer.

## Statistical analysis

Statistical analyses were performed in R (version 4.5.0). Comparisons between two groups were performed by two-sided Student’s *t*-test. For comparisons among multiple groups, one-way analysis of variance (ANOVA) followed by Dunnett’s test was performed using the DescTools package. Unless otherwise stated, statistical significance was set at *p* ≤ 0.05.

## Data and code availability

RNA-seq data are available through the Gene Expression Omnibus (GEO) under accession number GSE344432. Source code used for data visualization is available upon request.

## Supporting information

Supplementary Table 6: Target genes and sgRNAs in CRISPR library

Supplementary Data 2: DE analysis of RNA-seq

Supplementary Data 1: Raw count data for CRISPR screens

## Acknowledgement

We thank Uehara Memorial Foundation, International Medical Research Foundation, and The Osamu Hayaishi Memorial Scholarship for Study Abroad for supporting R.M. through a research fellowship.

## Author contributions

**R. Miyamoto:** Conceptualization, methodology, experiments, bioinformatic analysis, writing-original draft, writing-review and editing. **J. Park:** Experiments, methodology, writing-review and editing, **G. Dwosh**: Experiments. **V. Zuco:** Resources. **S. Pasquali:** Resources. **B. Dickson:** Histology. **D. Kirsch:** Funding acquisition, conceptualization, supervision, writing-review and editing.

---

**Supplementary Fig. 1.**
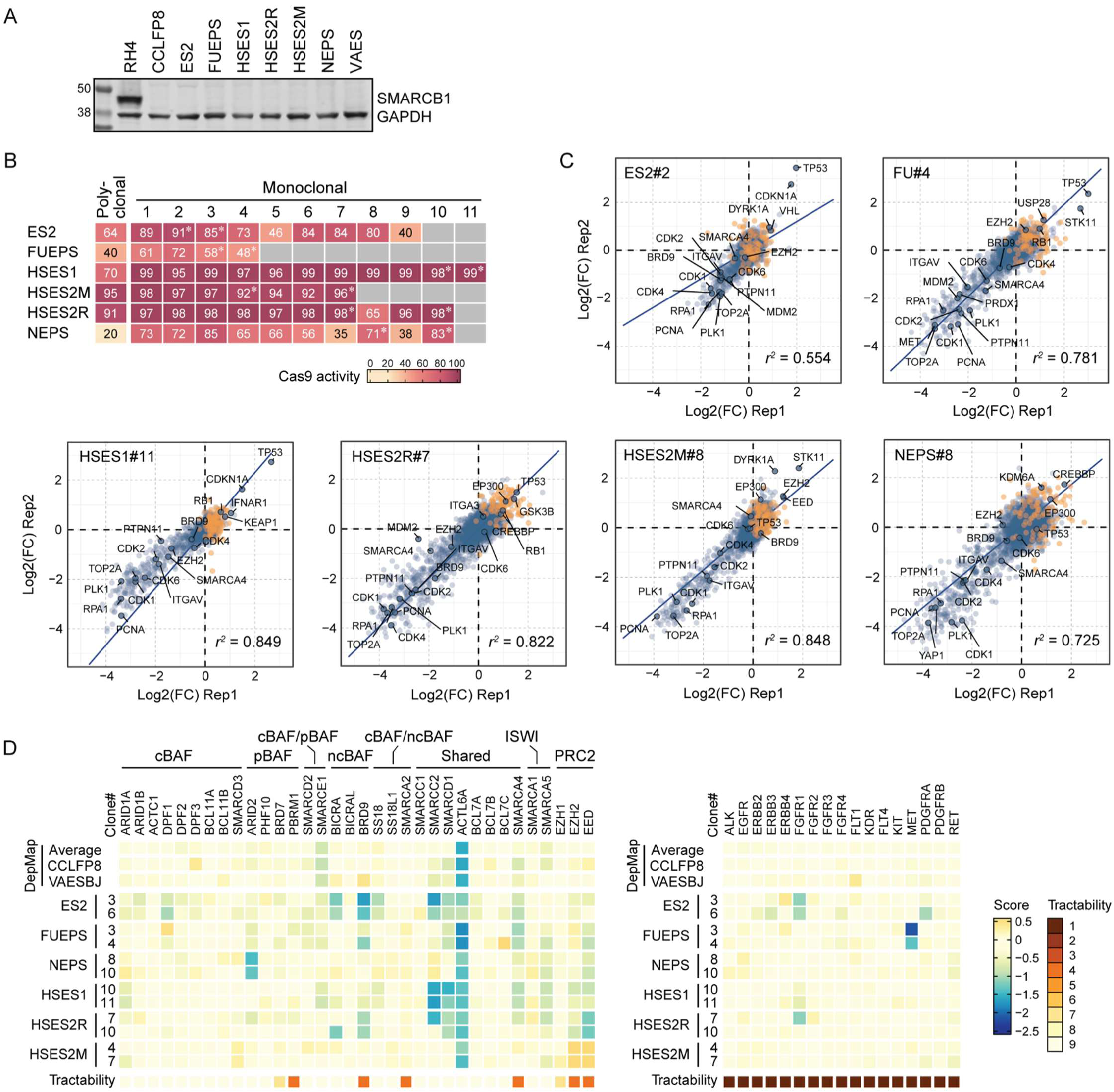
| Druggable gene-targeted CRISPR screens using preoptimized EPS cell models. (A) Western blotting showing loss of SMARCB1 in EPS cell lines used in this study. RH4 was used as a SMARCB1- positive sarcoma cell line. (B) Cas9 activity of polyclonal and monoclonal EPS cells expressing Cas9. Four to eleven monoclonal derivatives were established, and Cas9 activity was assessed based on editing efficacy at AAVS1 locus. *Clones used for screens. (C) Correlation of sgRNA depletion between two independent screen replicates. Well-characterized tumor suppressors, core essential genes, and genes investigated in this study are indicated. (D) Heatmaps showing dropout scores for (left) components of BAF, ISWI and PRC2 complexes, and (right) receptor tyrosine kinases with clinically available inhibitors.

**Supplementary Fig. 2.**
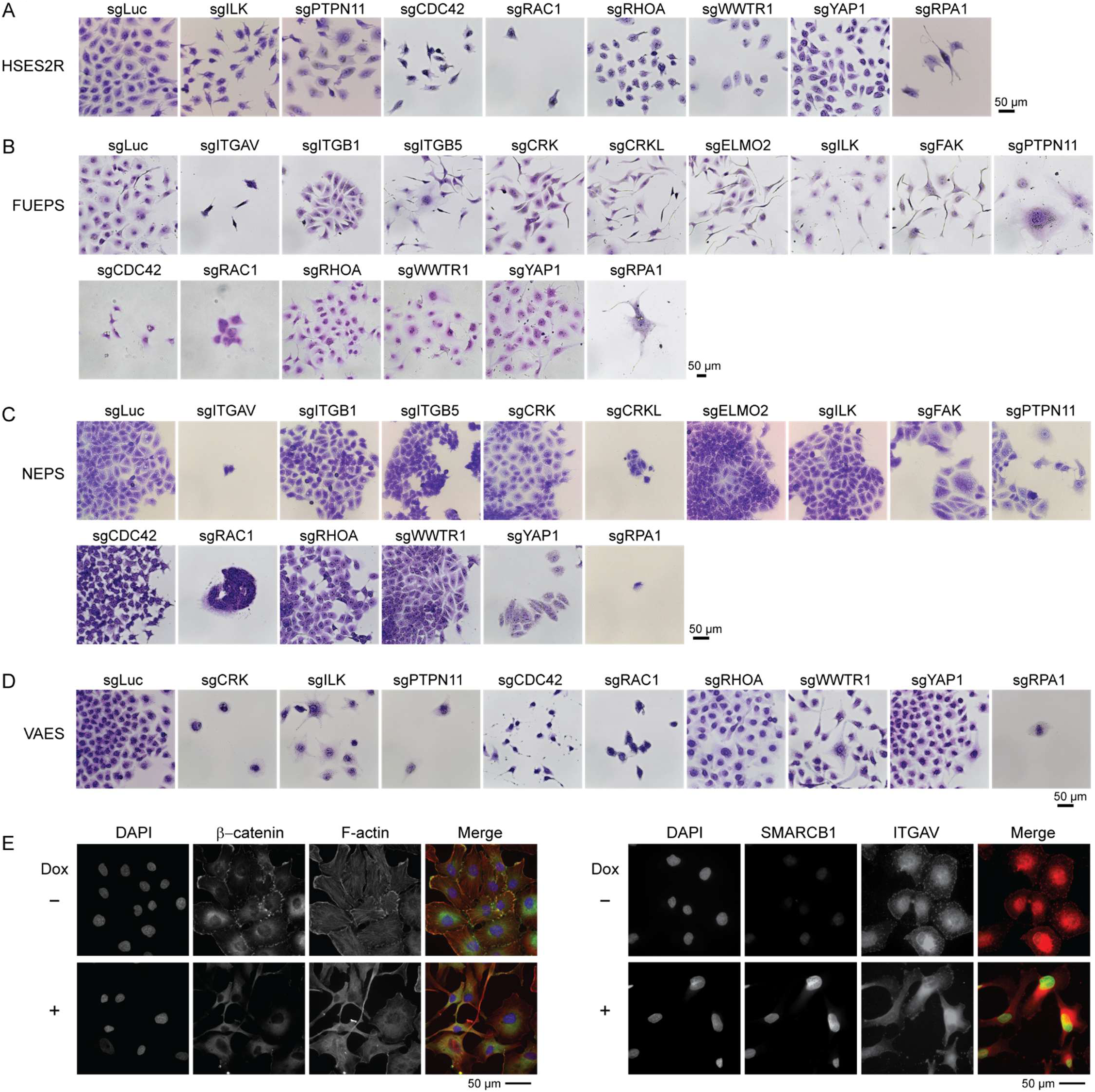
| Functional knockout of integrin signaling components disrupts epithelioid growth in EPS. (A-D) Morphological changes following CRISPR-mediated knockout of the indicated integrin signaling components in (A) HSES2R, (B) FUEPS, (C) NEPS, and (D) VAES cells. NEPS often retained epithelial morphology and cluster formation despite knockout of integrin signaling genes, highlighting biological heterogeneity in EPS. (E) Immunohistochemistry showing the distribution of (left) β-catenin and F-actin, and (right) SMARCB1 and ITGAV in FUEPS without or with SMARCB1 re-expression. A leaky SMARCB1 expression was detected in dox-untreated cells.

**Supplementary Fig. 3.**
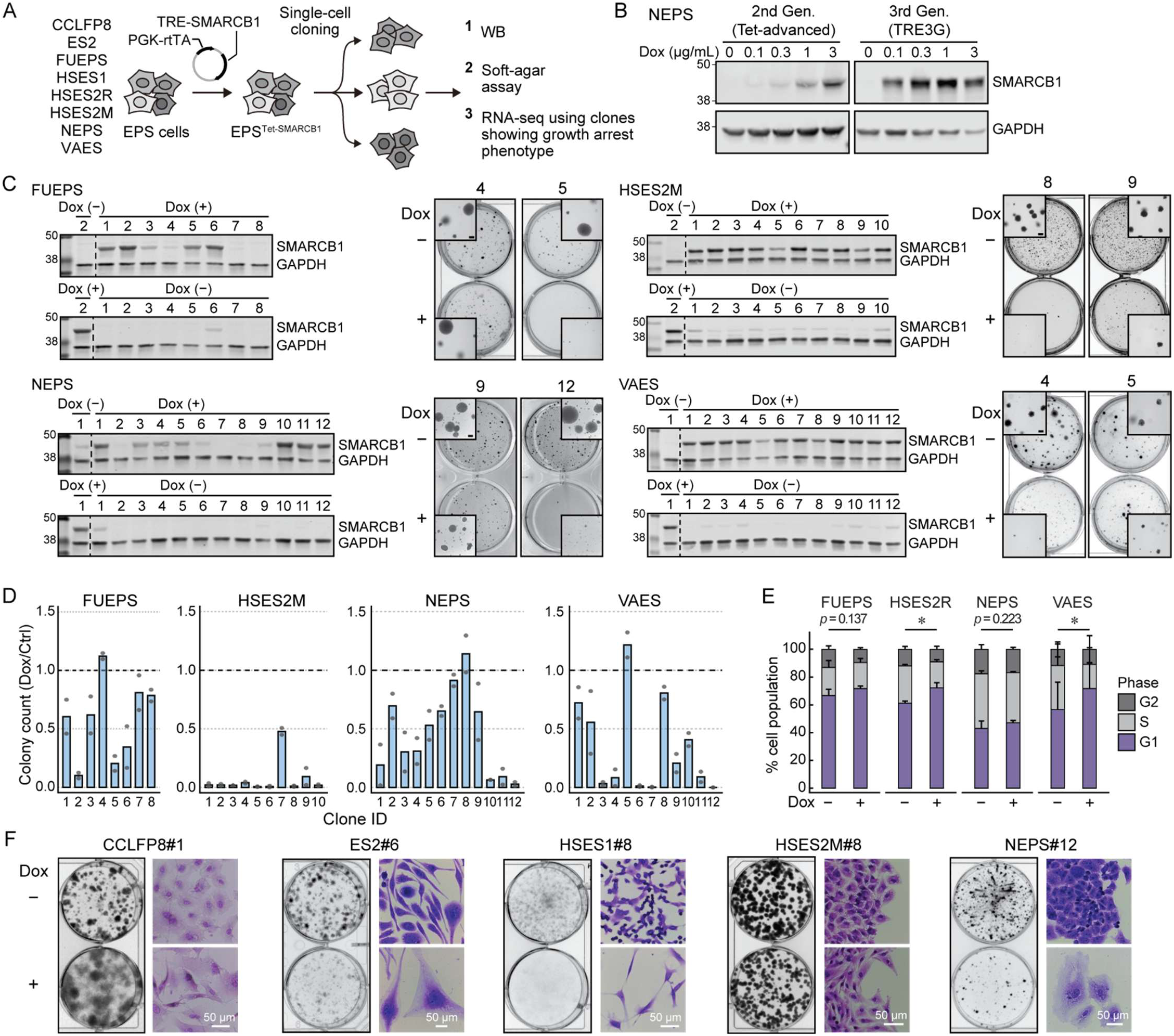
| SMARCB1 loss is a primary driver of epithelioid growth in EPS. (A) Schematic of subcloning strategy for SMARCB1 re-expression assays. (B) Comparison of SMARCB1 induction efficacy between second- and third-generation Tet-ON systems. Cells were treated with doxycycline (Dox) for 3 days at the indicated concentrations. (C) Clone-to-clone variation in Dox-induced SMARCB1 expression and corresponding effects on anchorage-independent colony formation. For protein expression analysis, cells were treated with 2 µg/mL Dox for 3 days. A reference sample was included in the first lane. Scale bars in the inset images indicate 200 µm. (D) Relative colony counts (Dox-treated/Dox-untreated) in soft agar assays. Bars represent the averages of two replicates, and dots represent individual replicates. (E) Cell-cycle analysis in EPS cells. Cells were stained with propidium iodide (PI) after 3-day culture with or without Dox. Values indicate mean ± SD. n = 3. Paired Student’s *t*-test was used to analyze the difference of G1 population. \**p* < 0.05. (F) Morphological changes following SMARCB1 re-expression in adherent clonogenic assay.

**Supplementary Fig. 4.**
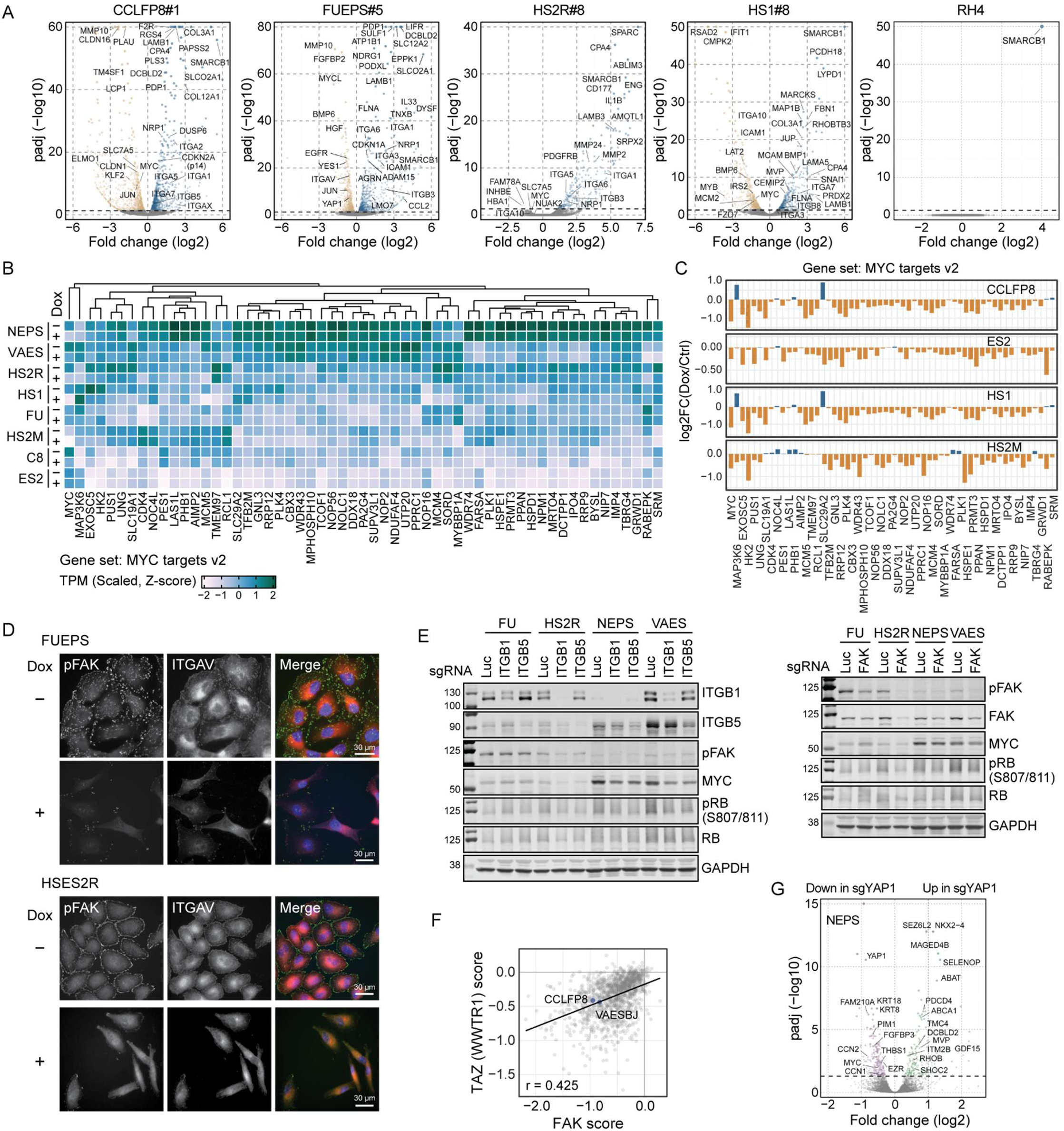
| SMARCB1-loss promotes integrin-YAP/TAZ-MYC signaling in EPS. (A) Volcano plots showing transcriptomic changes following SMARCB1 re-expression in EPS and an RMS cell line. Single-cell clones were used for EPS lines. (B) Expression levels of MYC target genes across eight EPS cell lines. TPM values were scaled across lines and are shown as Z-scores. Genes from the MSigDB MYC targets v2 are displayed. (C) Fold changes of MYC target genes in the indicated EPS lines. (D) ICC of FUEPS and HSES2R showing reduced or polarized pFAK expression following SMARCB1 restoration. (E) Reduced MYC and pRB levels in EPS cells following CRISPR knockout of ITGB1, ITGB5, and FAK. (F) Dependency score correlation between FAK and TAZ in DepMap. (G) Transcriptomic changes after YAP1 knockout in NEPS cells. Luciferase sgRNA was used as a control.

**Supplementary Fig. 5.**
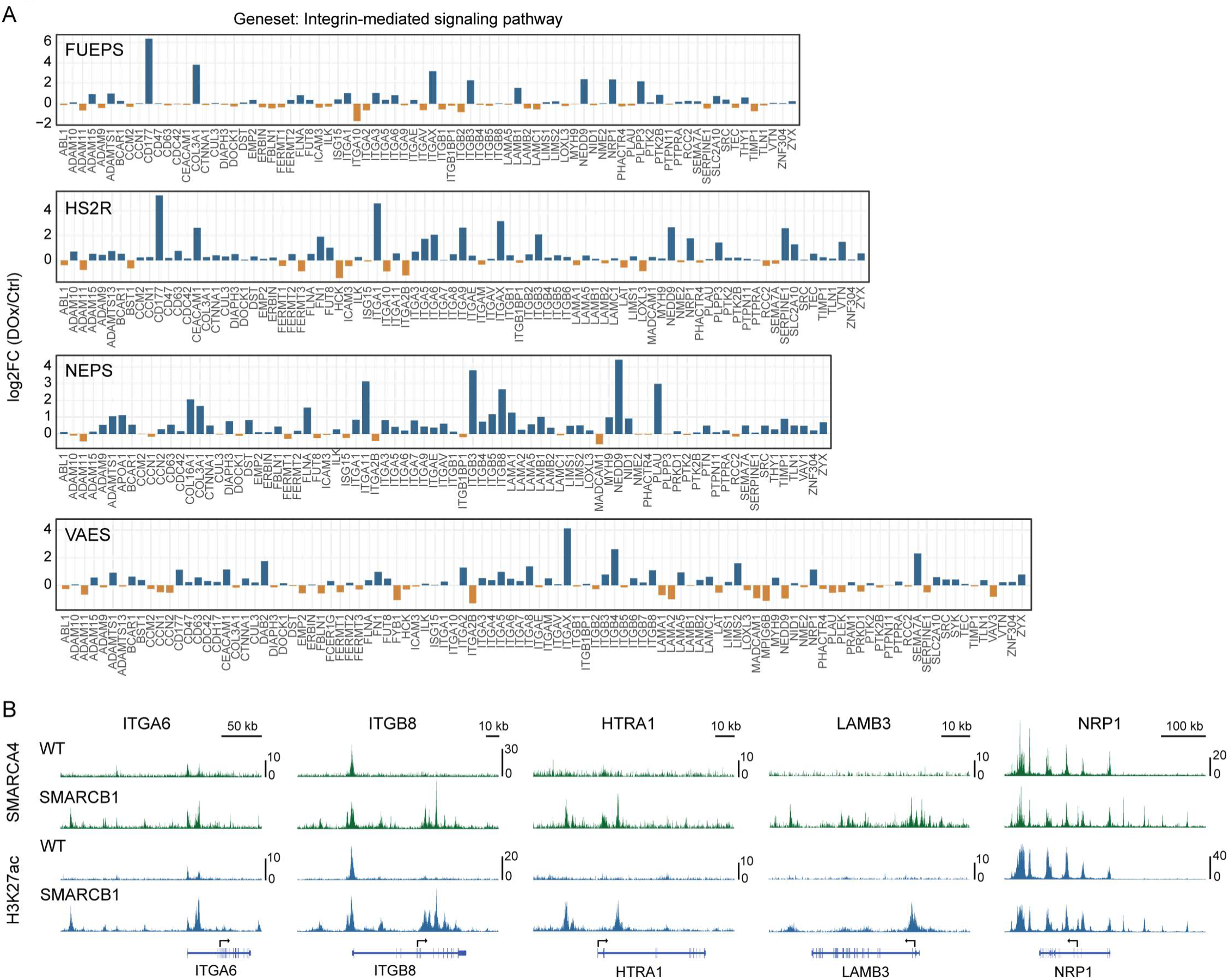
| Integrin signaling genes are direct targets of the BAF complex. (A) Expression changes of integrin signaling-associated genes after SMARCB1 re-expression. Genes from the MSigDB Integrin-mediated signaling pathway are shown. Genes expressed at low levels were omitted. (B) ChIP-seq profiles of SMARCA4 and H3K27ac in VAES cells. Peak tracks for Integrin receptor genes (ITGA6, ITGB8), the protease HTRA1, the laminin subunit LAMB3, and the integrin signaling modulator NRP1 are shown.

**Supplementary Fig. 6.**
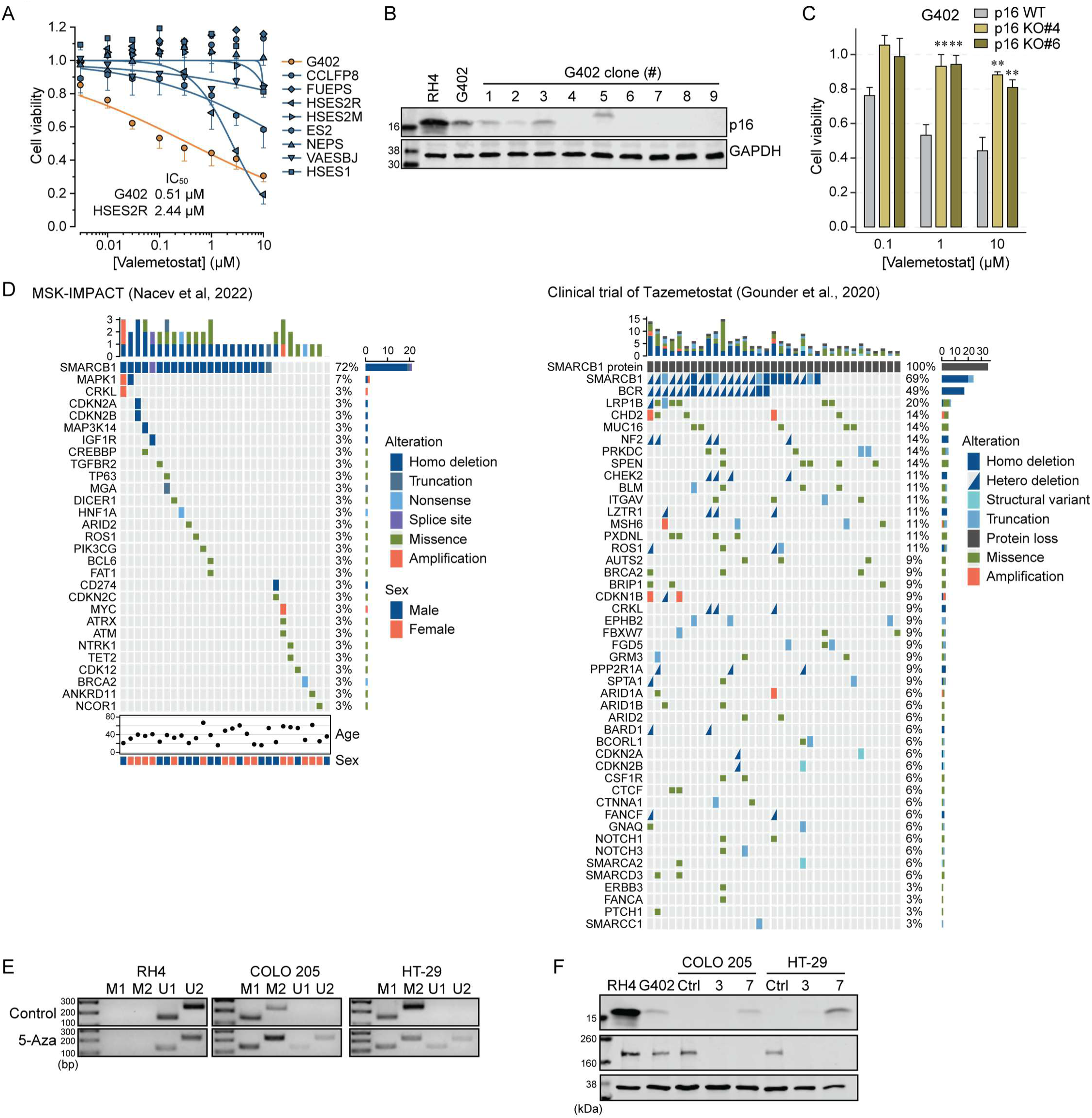
| CDKN2A alterations in clinical EPS tumors. (A) Cell viability assay for valemetostat. Cells were treated with valemetostat for 7 days. (B) p16 expression in nine G402-derived subclones subjected to p16 knockout procedures. To delete p16, Cas9 nickase and a pair of sgRNAs targeting CDKN2A exon 1α were transiently expressed. RH4 and parental G402 cells were used as controls. (C) Effect of p16 knockout on valemetostat-induced cytotoxicity in G402 cells. \*\**p* < 0.01. ANOVA followed by Dunnett’s test. (D) Mutational landscape of EPS. Data from two independent sources are summarized separately. (E) MS-PCR for RH4 (methylation-negative control) and two colon cancer cell lines, COLO 205 and HT-29 (methylation- positive control). (F) Recovery of p16 expression in COLO 205 and HT-29 after 5-Aza treatment for the indicated durations.

**Supplementary Fig. 7.**
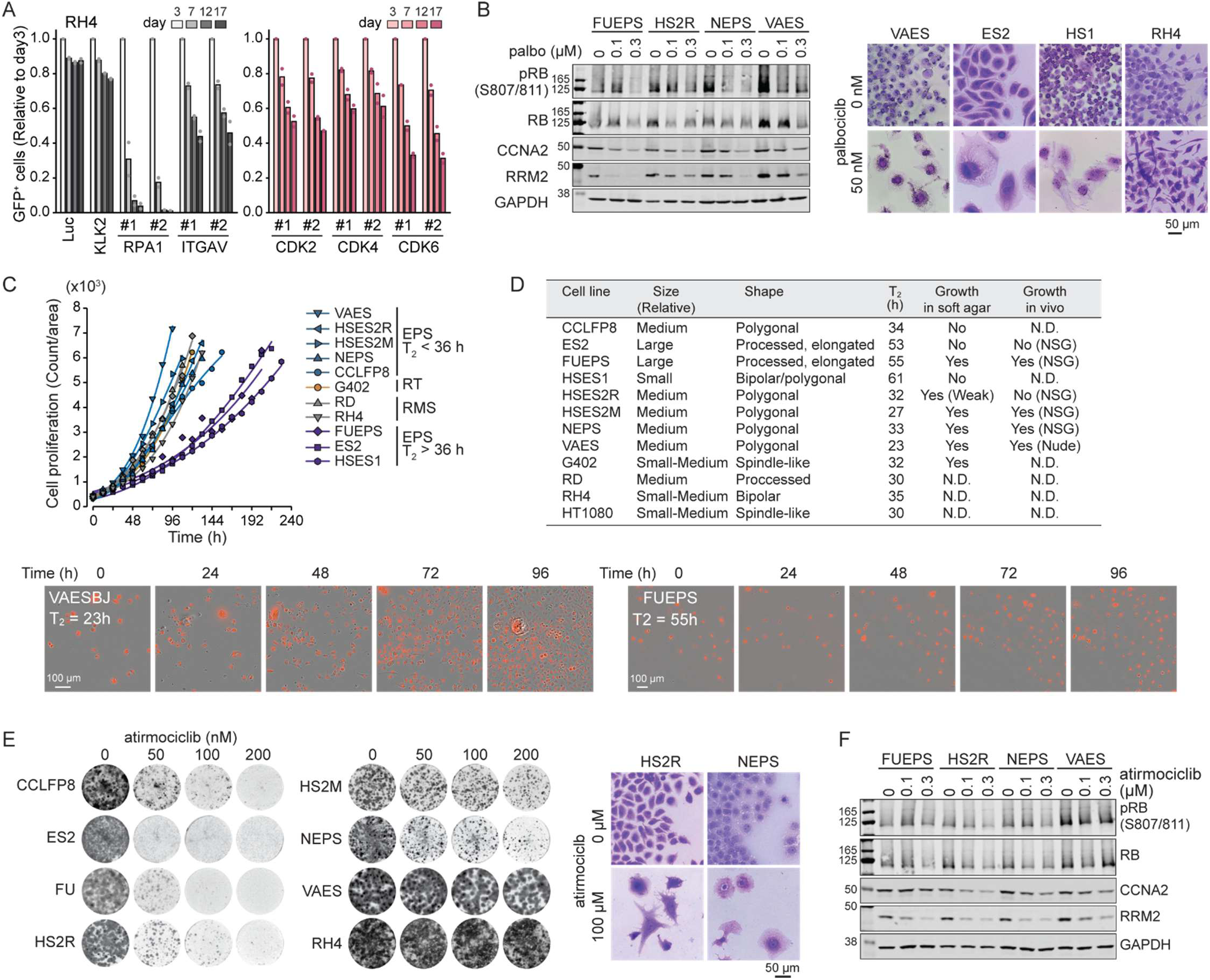
| CDK4/6 blockade in EPS. (A) CRISPR competitive growth assay in RH4 cells. (B) Effects of palbociclib on cell-cycle markers and morphology in EPS cells. Cells were treated with palbociclib for (left) 3 days or (right) 14 days. Scale bar indicates 50 µm. (C) Growth kinetics of cell lines used in this study. Cells stably expressing nuclei-localizing mCherry were analyzed using Incucyte. Cell count was determined based on nuclear red fluorescence. Representative Incucyte images of VAES and FUEPS are shown. T_2_, doubling time. (D) Cell line metadata based on observations in this study. N.D, not determined. (E) Selective growth-suppressive effects of atirmociclib in CDK4-dependent EPS cell lines. Cells were treated with atirmociclib for 14 days. (F) Effects of atirmociclib on cell-cycle markers in EPS cell lines. VAES, a CDK6-dependent cell line, showed a moderate phenotype compared with CDK4-dependent lines.

**Supplementary Fig. 8.**
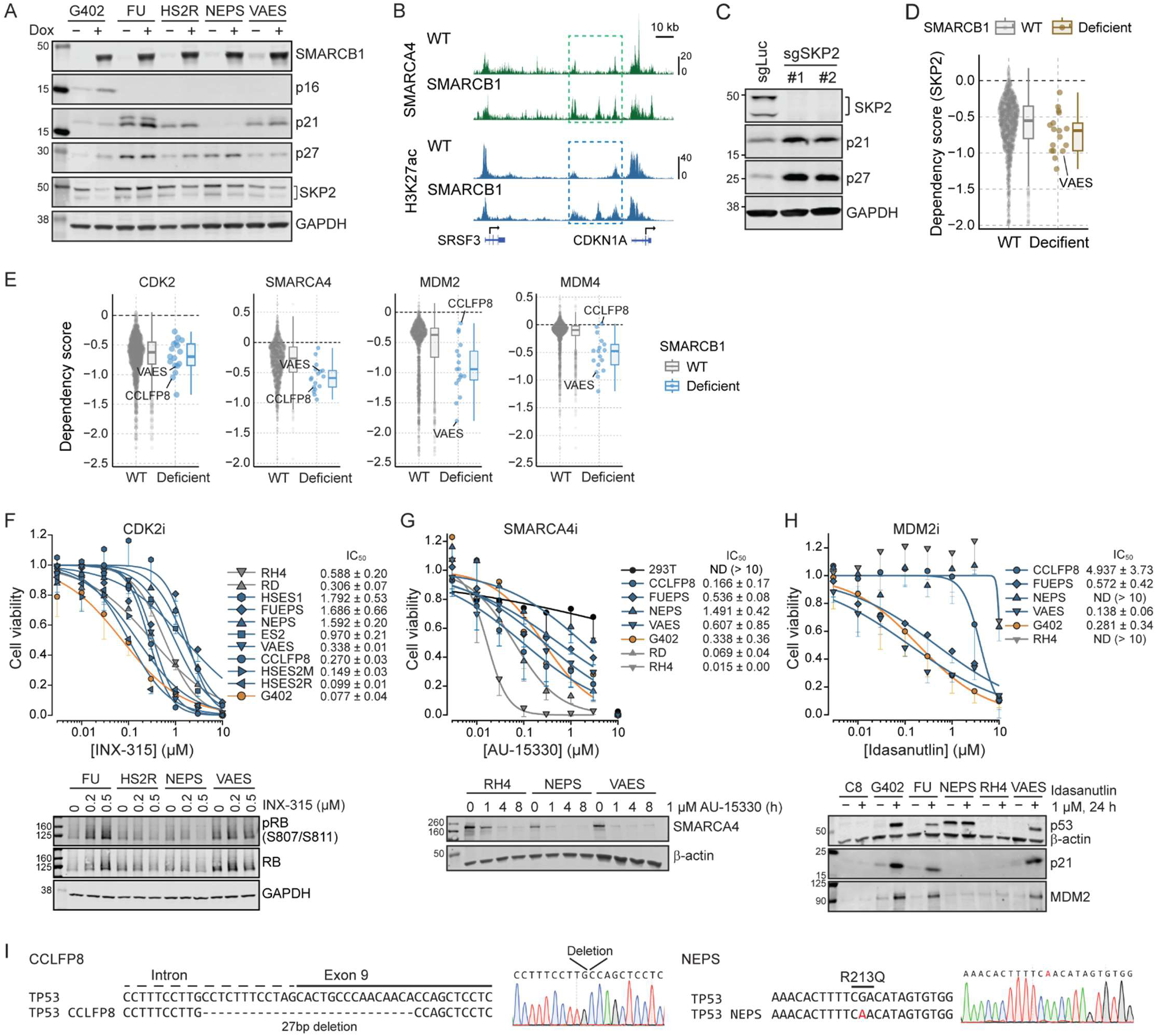
| Dysregulated cell-cycle control in EPS. (A) Expression of CDK inhibitor proteins (p16, p21, p27) and SKP2, a component of the SCF^SKP2^ complex, in EPS cells without and with SMARCB1 re-expression. Cell lysates were collected after 3-day culture without or with Dox. (B) ChIP-seq data showing occupancy of SMARCA4 and H3K27ac in VAES cells. SMARCB1 re-expression induced gain of SMARCA4 and H3K27ac peaks at a distal enhancer region of CDKN1A (p21). (C) CRISPR knockout of SKP2 and resultant upregulation of p21 and p27 in VAES cells. Cell lysates were collected 3 days after sgRNA transduction. (D) Two-class comparison of DepMap dependency scores between SMARCB1 wild-type and SMARCB1-deficient cancer cell lines. A subset of SMARCB1-deficient cell lines, including VAES, shows a significant SKP2 dependency. (E) Two-class comparison of dependency scores for CDK2, SMARCA4, MDM2, and MDM4. (F-H) Cell viability assay for (F) the CDK2 inhibitor INX-315, (G) the SMARCA4 inhibitor AU-15330, and (H) the MDM2 inhibitor idasanutlin. n = 3, mean ± S.D. Western blotting data show (F) the effect of INX-315 on RB/pRB levels, (G) SMARCA4 degradation following AU-15330 treatment, and (H) induction of p53 and its target proteins p21 and MDM2 following idasanutlin treatment. INX-315-induced elevation of pRB in FUEPS suggests a presence of compensatory mechanisms. (I) Sequencing analysis of TP53 exons in CCLFP8 and NEPS cells. Biallelic small deletions or mutation (R213Q) was identified, suggesting a potential resistance mechanism of MDM2 inhibition.

**Supplementary Fig. 9.**
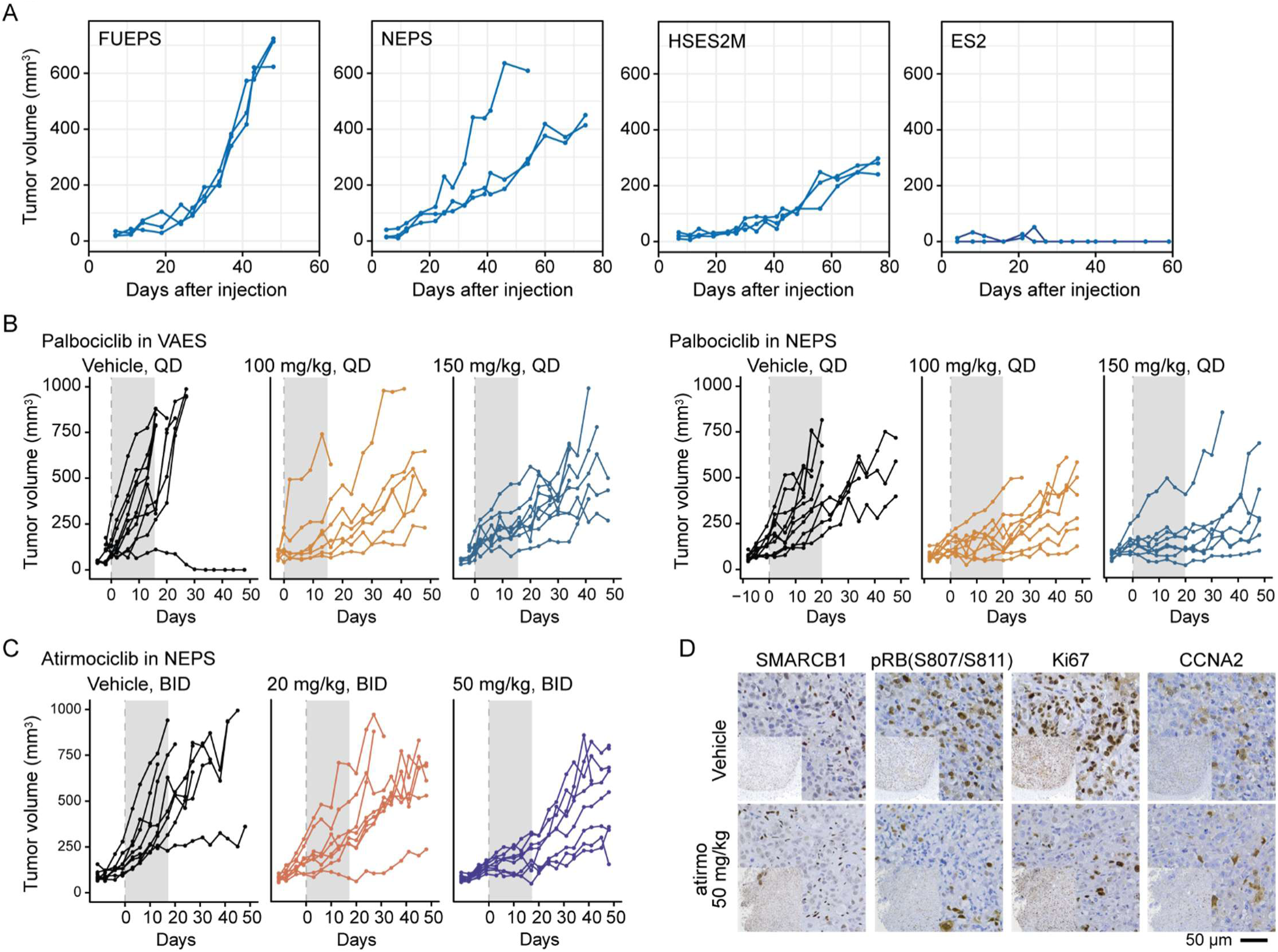
| Kinetics of EPS cell growth *in vivo* (A) Tumor forming capacity of EPS cell lines *in vivo*. Five million cells from the indicated cell lines were implanted subcutaneously in the flanks of NSG mice, and the tumor growth was monitored for up to 80 days with weekly tumor measurements. (B, C) Growth curves of individual tumors in *in vivo* testing of (B) palbociclib and (B) atirmociclib. Grey shadowing indicates drug treatment. Q.D, once daily. BID, twice daily. (D) IHC of proliferation markers in vehicle-or atirmociclib-treated NEPS tumors.

**Supplementary Table 1.**
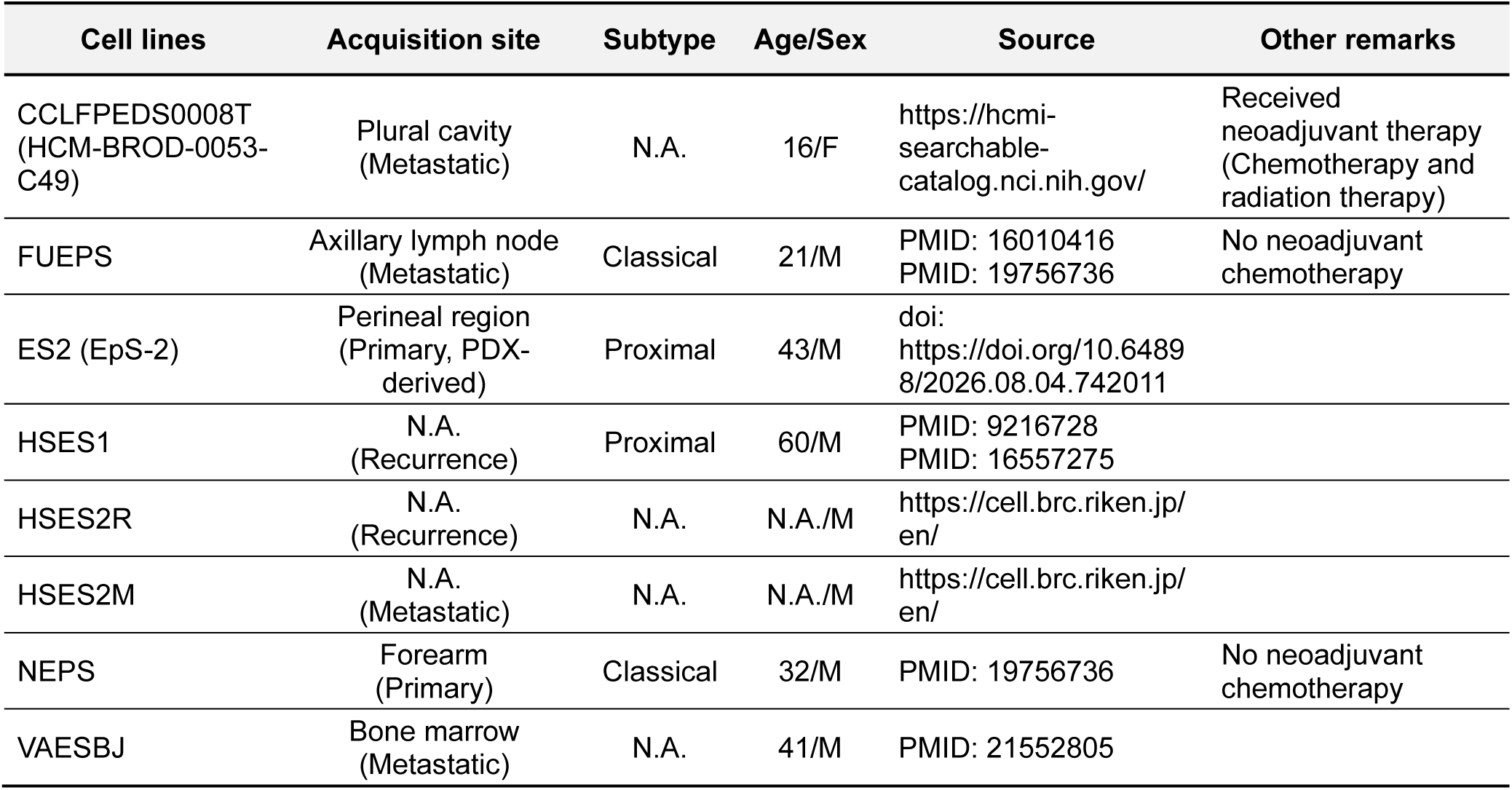
List of cell lines

**Supplementary Table 2.** p16 IHC scores and sample information

| Name | Percentage | Intensity | Modified Allred score | Original diagnosis | Anatomic site | Primary vs Metastasis | Age | Sex |
| --- | --- | --- | --- | --- | --- | --- | --- | --- |
| Untreated |  |  |  |  |  |  |  |  |
| ES-1 | 0 | 0 | 0 | Epithelioid sarcoma | Arm | Metastasis | 32 | Female |
| ES-2 | 0 | 0 | 0 | Epithelioid sarcoma | Toe | Primary | 30 | Female |
| ES-3 | 3 | 3 | 6 | Epithelioid sarcoma | Abdominal wall | Primary | 47 | Male |
| ES-4 | 4 | 1 | 5 | Epithelioid sarcoma (SMARCB1-deficient vulvar neoplasm) | Groin | Primary | 58 | Female |
| ES-5 | 0 | 0 | 0 | Epithelioid sarcoma | Intra-abdominal | Metastasis | 38 | Male |
| ES-6 | 0 | 0 | 0 | Epithelioid sarcoma | Leg | Unknown | 80 | Male |
| ES-7 | 3 | 2 | 5 | Epithelioid sarcoma | Foot | Primary | 29 | Male |
| ES-8 | 0 | 0 | 0 | Epithelioid sarcoma | Calf | Primary | 69 | Female |
| ES-9 | 0 | 0 | 0 | Epithelioid sarcoma (SMARCB1-deficient vulvar neoplasm) | Vulva | Primary | 49 | Female |
| With neoadjuvant therapy |  |  |  |  |  |  |  |  |
| ES-10 | 0 | 0 | 0 | Epithelioid sarcoma (SMARCB1-deficient vulvar neoplasm) | Mons pubis | Primary | 51 | Female |
| ES-11 | 5 | 3 | 8 | Epithelioid sarcoma | Groin | Primary | 42 | Female |
| ES-12 | 5 | 3 | 8 | Epithelioid sarcoma | Inguinal | Primary | 47 | Male |
| ES-13 | 0 | 0 | 0 | Epithelioid sarcoma | Breast | Metastasis | 22 | Female |
| ES-14 | 0 | 0 | 0 | Epithelioid sarcoma | Forearm | Primary | 30 | Female |
| ES-15 | 4 | 2 | 6 | Epithelioid sarcoma (SMARCB1-deficient vulvar neoplasm) | Mons pubis | Primary | 50 | Female |

Allred score (Allred DC *et al. Mod Pathol* 1998;11:155–68)
| % Staining score | Proportion of positive staining | Intensity Score | Average intensity of positively stained cells |
| --- | --- | --- | --- |
| 0 | None | 0 | None |
| 1 | < 1/100 | 1 | Weak |
| 2 | 1/100 to 1/10 | 2 | Intermediate |
| 3 | 1/10 to 1/3 | 3 | Strong |
| 4 | 1/2 to 2/3 |  |  |
| 5 | > 2/3 |  |  |

**Supplementary Table 3.** List of antibodies

Antibodies for western blotting
| Target | Source | Identifier |
| --- | --- | --- |
| SMARCB1 | Cell Signaling Technology | 91735 |
| pRB (S807/811) | Cell Signaling Technology | 8516 |
| RB | Cell Signaling Technology | 9309 |
| CCNA2 | Cell Signaling Technology | 67955 |
| RRM2 | Cell Signaling Technology | 65939 |
| GAPDH | Proteintech | 60004-1-Ig |
| FAK | Cell Signaling Technology | 71433 |
| pFAK (Y397) | Cell Signaling Technology | 8556 |
| pFAK (Y925) | Cell Signaling Technology | 3284 |
| ITGAV | Abcam | ab179475 |
| MYC | Cell Signaling Technology | 13987 |
| TAZ | Proteintech | 23306-1-AP |
| YAP1 | Cell Signaling Technology | 14074 |
| ITGB1 | Abcam | ab134179 |
| ITGB5 | Abcam | ab309092 |
| Tubulin | Cell Signaling Technology | 86298 |
| ITGB3 | Abcam | ab119992 |
| ITGB8 | Proteintech | 29775-1-AP |
| ITGA1 | Abcam | ab317553 |
| ITGA2 | Abcam | ab181548 |
| ITGA3 | Proteintech | 21992-1-AP |
| ITGA5 | Abcam | ab150361 |
| ITGA6 | Proteintech | 27189-1-AP |
| p16 | Abcam | ab-108349 |
| p14 | Cell Signaling Technology | 74560 |
| H3K27me3 | MilliporeSigma | 07-449 |
| DNMT1 | Cell Signaling Technology | 5032 |
| CDK4 | Cell Signaling Technology | 12790 |
| CDK6 | Cell Signaling Technology | 13331 |
| Actin | Proteintech | 66009-1-Ig |
| p21 | Cell Signaling Technology | 2947 |
| p27 | Cell Signaling Technology | 3686 |
| SKP2 | Proteintech | 15010-1-AP |
| SMARCA4 | Abcam | ab110641 |
| p53 | Cell Signaling Technology | 2524 |
| MDM2 | Cell Signaling Technology | 86934 |

Antibodies for immunocytochemistry
| Target | Source | Identifier |
| --- | --- | --- |
| b-catenin | Cell Signaling Technology | 8480 |
| ZO-1 | Proteintech | 21773-1-AP |
| Phalloidin | Proteintech | PF00003 |
| ITGAV | Santa Cruz Biotechnology | sc-9969 |

Antibodies for immunohistochemistry
| Target | Source | Identifier |
| --- | --- | --- |
| SMARCB1* | BD | 612111 |
| SMARCB1 | Cell Signaling Technology | 91735 |
| p16* | Dako | GA783 |
| Ki-67 | Cell Signaling Technology | 12202 |
| CDK6 | Abcam | ab124821 |

\*For primary EPS samples

**Supplementary Table 4.**
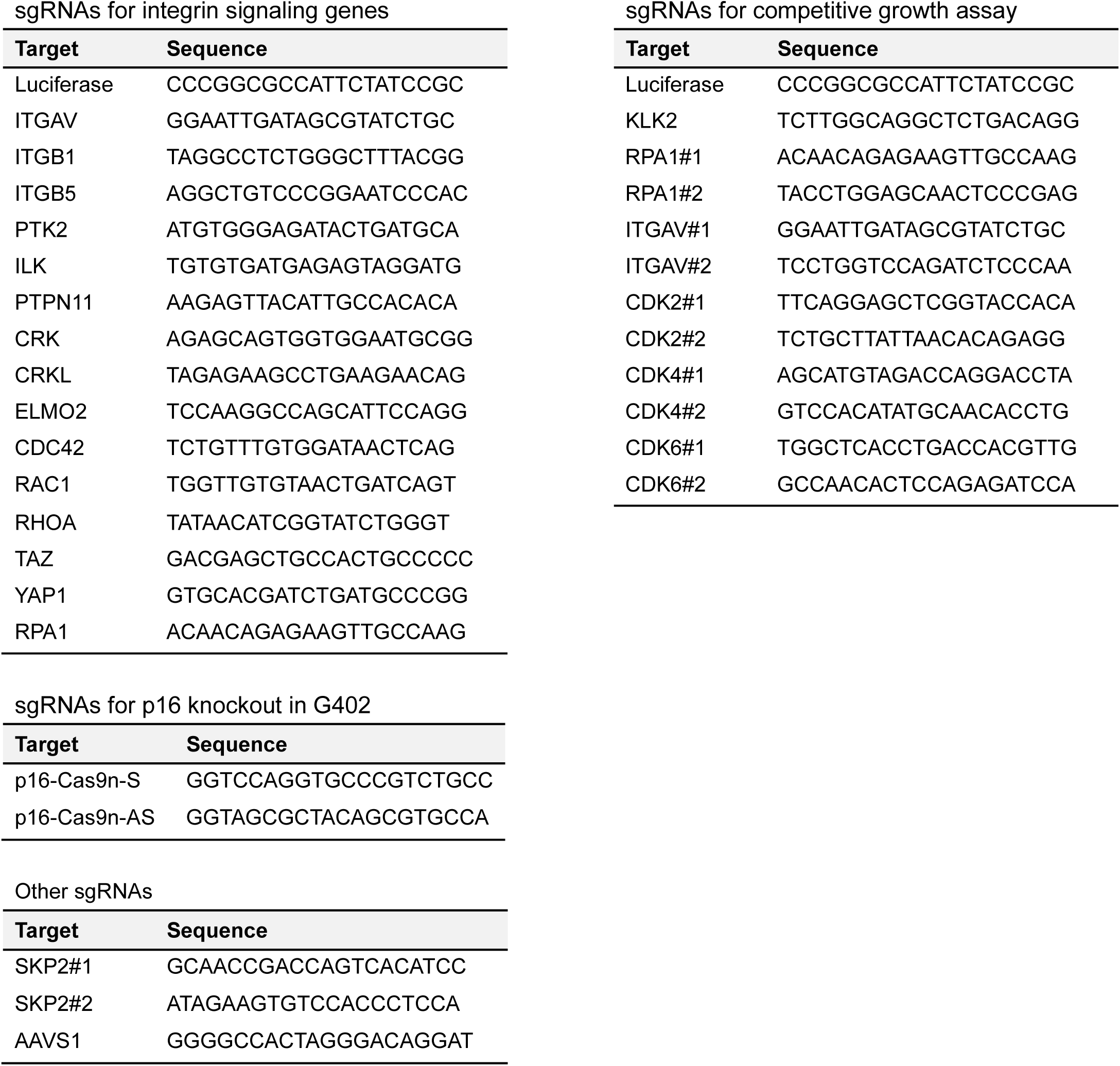
List of sgRNAs

**Supplementary Table 5.**
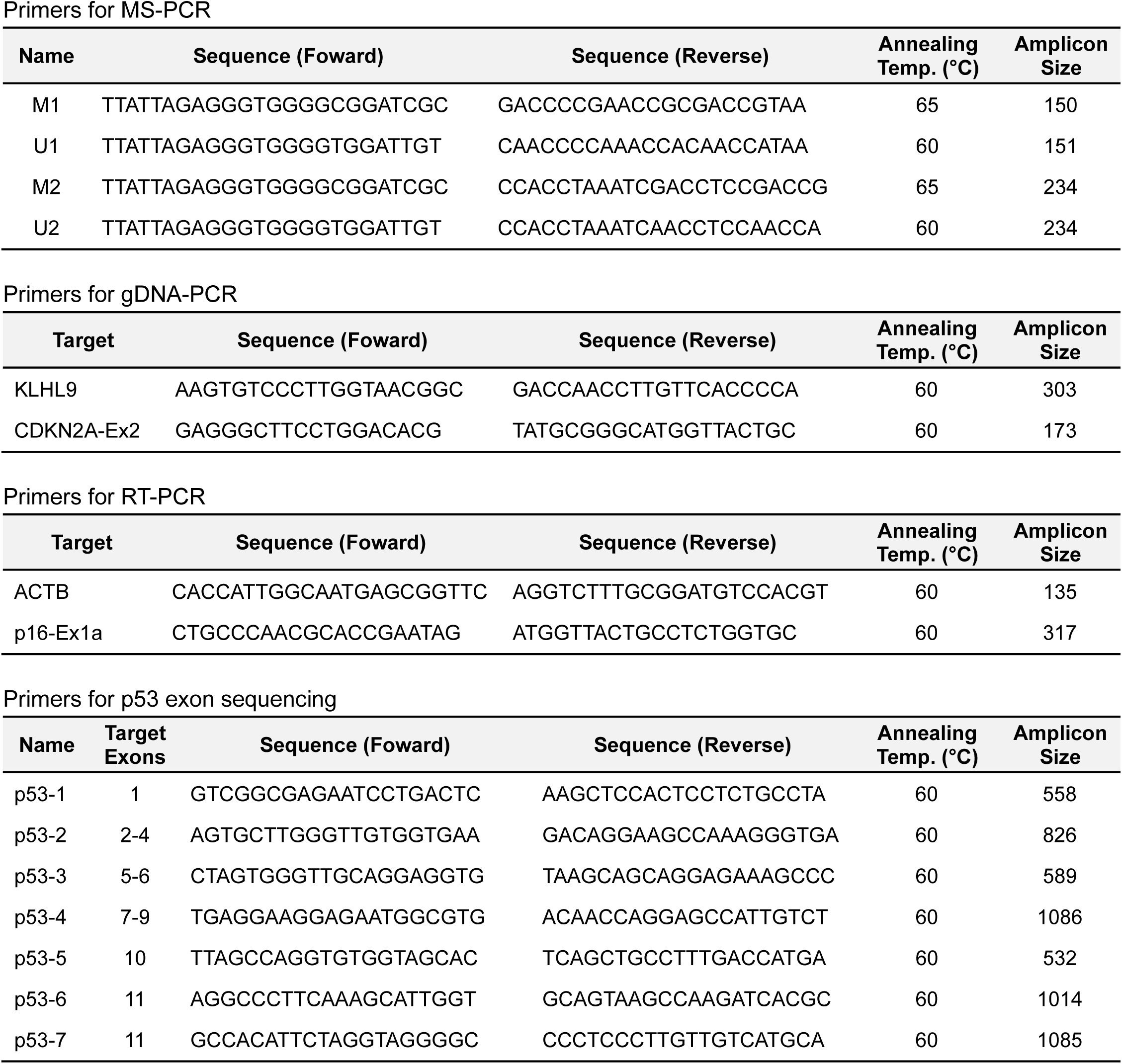
List of PCR primers

